# The receptor PTPRU is a redox sensitive pseudophosphatase

**DOI:** 10.1101/805119

**Authors:** Iain M. Hay, Gareth W. Fearnley, Pablo Rios, Maja Köhn, Hayley J. Sharpe, Janet E. Deane

## Abstract

The dynamic regulation of protein tyrosine phosphorylation is a critical feature of intercellular communication and is regulated by the actions of kinases and phosphatases. The receptor-linked protein tyrosine phosphatases (RPTPs) are key signaling molecules that possess an extracellular domain and intracellular phosphatase domains. Most human RPTPs have tandem intracellular tyrosine phosphatase domains: a catalytically active membrane proximal (D1) domain; and a membrane distal (D2) inactive “pseudophosphatase” domain. The receptor PTPRU plays a role in development, multiple cancers and has been implicated in the dephosphorylation of cell adhesion proteins. However, PTPRU has a non-canonical D1 domain containing several sequence variations in key catalytic loops that suggest it may function using a mechanism distinct from related RPTPs. Here, we demonstrate through biochemical and structural studies that PTPRU is unique amongst the RPTPs in possessing two pseudophosphatase domains. We show that PTPRU-D1 displays no detectable catalytic activity against a range of phosphorylated substrates and determine that this is due to substantial disorder in the substrate-binding pocket as well as rearrangement of the catalytic loop such that the active site cysteine is occluded. We also show that this cysteine can form an intramolecular disulfide bond with a vicinal “backdoor” cysteine. Further, we demonstrate that the PTPRU D2 domain can recruit substrates of related RPTPs suggesting that this pseudophosphatase functions by competing with active phosphatases for the binding of substrates involved in cell adhesion. Therefore, PTPRU is a *bona-fide* pseudophosphatase and its functional role in cell signaling is via a non-catalytic mechanism.

**SIGNIFICANCE STATEMENT:** Protein tyrosine phosphorylation is a key post-translational modification required for cellular communication that is dynamically regulated by the activities of tyrosine kinases and phosphatases. Receptor tyrosine phosphatases (RPTPs) possess an extracellular receptor domain and intracellular phosphatase domains. We show that PTPRU is a non-canonical RPTP devoid of catalytic activity and demonstrate that this is due to multiple structural rearrangements. Despite this, PTPRU retains the capacity to bind the substrates of related phosphatases suggesting that the non-catalytic function of this pseudophosphatase is to compete with active phosphatases for the binding of substrates. Such pseudoenzymes represent an exciting and growing area of research with implications as key regulators of signaling networks.

## INTRODUCTION

The human classical protein tyrosine phosphatases (PTPs) are key signaling regulators that work with kinases to fine-tune cellular levels of phosphotyrosine, impacting on multiple cellular pathways including metabolism, differentiation, and adhesion (1–3). PTPs do not simply function as negative regulators of tyrosine kinases to reverse protein phosphorylation, instead it is becoming clear that PTPs work in synergy with kinases to regulate complex cell signaling pathways and are important therapeutic targets in diseases such as cancer and diabetes (4, 5). The 37 classical PTPs exhibit diverse domain architectures and subcellular localizations, but all share a conserved core catalytic C-X_5_-R motif, known as the PTP loop, which includes the essential cysteine that catalyzes the nucleophilic attack on the substrate phosphate group (6, 7). Less well conserved are the three additional motifs that form the PTP active site (7). The WPD loop contains an aspartate residue that acts as a general acid/base during different steps of the catalytic cycle and assists in substrate binding. The phosphotyrosine (pTyr) recognition loop, typically containing the protein sequence KNRY, is so called because it forms the deep pocket that imparts selectivity for phosphotyrosine over smaller phosphorylated amino acids, such as serine and threonine. In addition to defining the shape of the binding pocket, the tyrosine in the pTyr recognition loop plays a crucial role in substrate orientation as its sidechain packs against the substrate phosphotyrosine phenyl ring. Finally, the Q loop positions and activates a water molecule for the hydrolysis of the phosphocysteine intermediate complex.

Despite the importance of catalysis for the function of many PTPs, there are increasing reports of non-catalytic functions (8–10). Moreover, 5 of the 37 classical PTPs are reported to be catalytically inactive, including the non-receptor PTPs: PTPN23 (HD-PTP) (11, 12), PTPN14 (PTPD2) and PTPN21 (PTPD1) (11, 13) and the receptor PTPs: PTPRN (PTPIA2) and PTPRN2 (PTPIA2β) (14). These pseudoPTPs frequently contain altered sequences in their catalytic motifs and substrate binding loops. For example, PTPN23 (HDPTP) has an incomplete Q loop, PTPN21 (PTPD1) possesses an altered WPD motif and PTPN14 (PTPD2) has a variant pTyr recognition loop (11, 13). However, the absence of enzyme function does not diminish the importance of these proteins in cell function, for example, PTPN23 is encoded by an essential gene (15, 16). Furthermore, 12 of the 21 cell surface receptor PTPs possess highly conserved membrane distal pseudophosphatase domains, which have been implicated in substrate recognition, redox sensing and enzyme regulation (3, 17, 18). Interestingly, changes in the catalytic loops do not always result in loss of catalytic activity but instead can alter the substrate specificity. For example, a glutamate substitution in the WPD motif of the single PTP domain of PTPRQ (PTPS31) determines its selectivity for phosphoinositides over phosphotyrosine (19).

PTPRU is a member of the R2B receptor family, which includes PTPRK, PTPRM and PTPRT. PTPRU is expressed during development (20, 21), and is reported to function during zebrafish gastrulation (22) and chick midbrain development (23). Interestingly, while PTPRK and PTPRT are reportedly tumor suppressors, PTPRU has been proposed to play an oncogenic role in gastric cancer and glioma cells (24, 25). The R2B receptors are characterized by a large extracellular domain that mediates homophilic interactions and tandem intracellular phosphatase domains (**Fig. 1A**) (26). The membrane proximal (D1) domains of PTPRK, PTPRM and PTPRT are active tyrosine phosphatases (11), however, the catalytic activity of PTPRU has not been determined. PTPRU possesses non-canonical WPD (WPE) and pTyr recognition loop sequences (GSRQ rather than KNRY) as well as a unique threonine in the PTP loop (**Fig. 1B** and **SI Appendix, Fig. S1**). Despite these sequence variations, previous work suggests that PTPRU can dephosphorylate substrates such as β-catenin (27, 28). Recently, it was shown that Afadin, a PTPRK substrate (3), is an interactor of PTPRU (29), implicating it as a cell adhesion regulator. To better understand the function of PTPRU we set out to determine whether it is an active phosphatase.

**Fig. 1.**
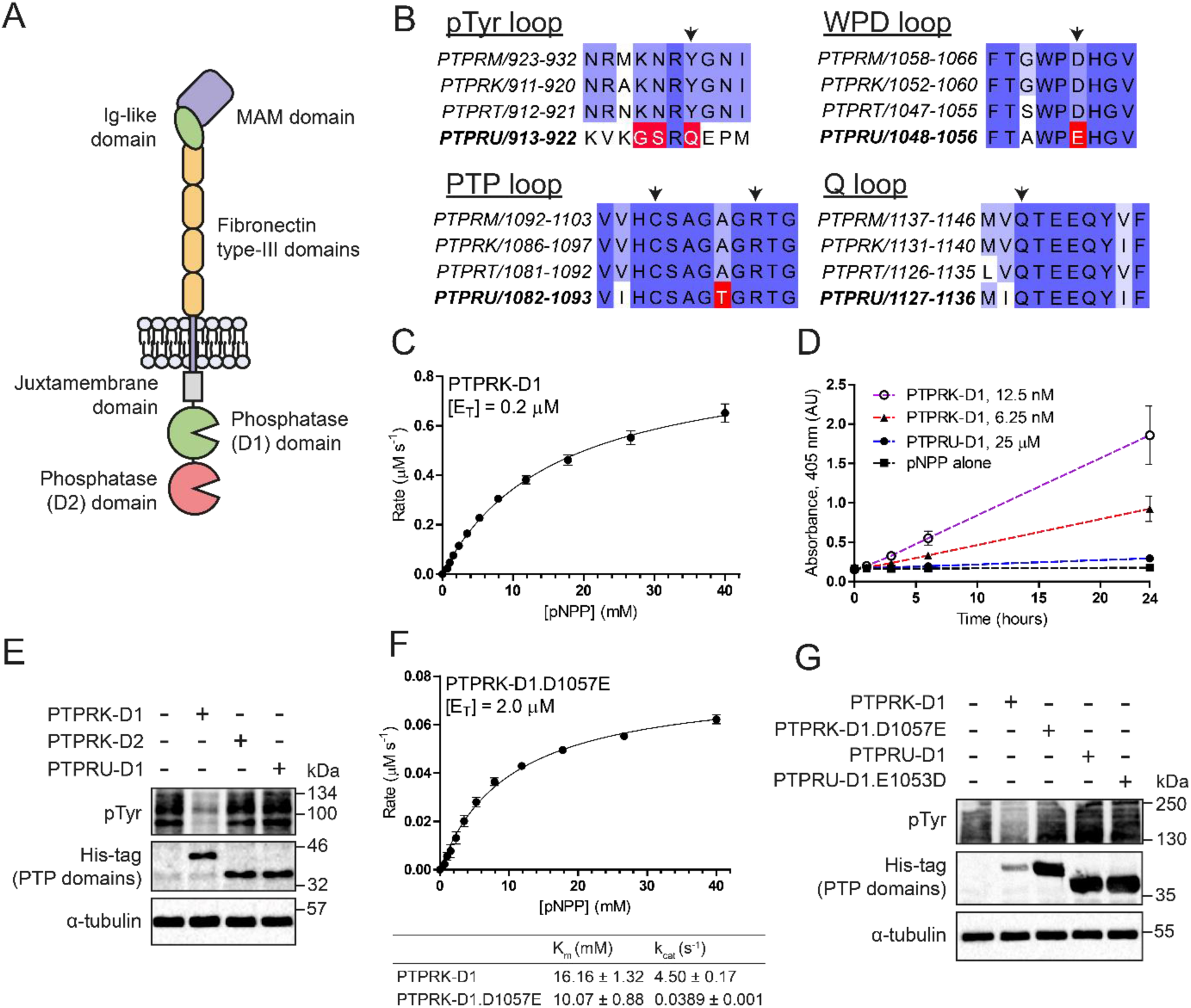
The PTPRU D1 domain does not dephosphorylate pTyr. *(A)* Schematic diagram of the R2B family RPTP domain structure. *(B)* Multiple sequence alignment of the 4 key PTP motifs across R2B family RPTPs, colored by percentage identity (blue). Key variable residues in PTPRU are highlighted in red and essential PTP catalytic residues are marked by arrowheads. *(C)* Michaelis-Menten plot of initial rate vs. substrate (pNPP) concentration using 0.2 μM PTPRK-D1. Error bars represent ± SEM of *n* = 3. *(D)* Extended time course of pNPP dephosphorylation, monitored by absorbance at 405 nm, using low concentration (6.25 and 12.5 nM) PTPRK-D1 and high concentration (25 μM) PTPRU-D1 recombinant proteins. *(E)* Immunoblot analysis of pervanadate-treated MCF10A lysates incubated with 0.1 μM PTPRK-D1, PTPRK-D2 or PTPRU-D1 recombinant PTP domains for 16 h at 4°C. *(F)* Michaelis-Menten plot of initial rate vs. substrate (pNPP) concentration using 2.0 μM PTPRK-D1.D1057E. Error bars represent ± SEM of *n* = 3. Table shows kinetic parameters K_m_ and k_cat_ ± SEM. *(G)* Immunoblot analysis of pervanadate-treated MCF10A lysates incubated with PTPRK-D1 (0.1 μM), PTPRK-D1.D1057E, PTPRU-D1 or PTPRU-D1.E1053D (5 μM) recombinant PTP domains for 16 h at 4°C.

Here, we demonstrate through biochemical and structural studies that PTPRU is unique amongst the RPTPs in possessing two pseudophosphatase domains. The crystal structure of the PTPRU D1 domain reveals substantial structural rearrangements to key catalytic loops such that the shape of the substrate binding pocket is lost and the active site cysteine is occluded. Despite lacking catalytic activity, PTPRU can recruit substrates of catalytically active paralogs supporting a model where PTPRU functions as a scaffold to compete for binding to protein substrates. Thus, the balance of homophilic RPTPs expressed on the surface of a cell will determine the local phosphotyrosine level of a subset of cell junction regulators.

## RESULTS

### PTPRU is catalytically inactive

To determine the consequence of sequence variations on PTPRU phosphatase activity, we expressed and purified recombinant D1 domains of PTPRU and the related PTPRK in *E. coli* for use in *in vitro* phosphatase assays. The generic substrate 4-nitrophenyl phosphate (pNPP) was used in initial activity assays. The K_m_ and k_cat_ for PTPRK-D1-mediated pNPP hydrolysis were 16.16 ± 1.32 mM and 4.50 ± 0.17 s^-1^, respectively, similar to previously determined kinetic parameters for the prototypic phosphatase PTP1B using this substrate (**Fig. 1C**) (30). Strikingly, we were unable to detect activity for PTPRU-D1 against pNPP, even at high enzyme concentrations (up to 25 μM) or when using an extended assay duration (**Fig. 1D**), whereas PTPRK-D1 activity was readily detectable in the low nanomolar range (**Fig. 1D**). To investigate whether a cellular cofactor might be necessary for PTPRU activity, we performed dephosphorylation assays using quenched pervanadate-treated cell lysates (see SI materials and methods), enriched in tyrosine phosphorylated proteins, as the substrate for recombinant PTP domains. Again, whilst incubation with PTPRK-D1 for 16 h at 4°C resulted in visible dephosphorylation of total cellular pTyr, PTPRU-D1 showed no activity and is comparable to the inactive PTPRK D2 domain (**Fig. 1E**). Taken together, these data suggest that unlike PTPRK, the D1 domain of PTPRU has no intrinsic PTP activity.

Due to the highly divergent nature of the PTPRU pTyr recognition loop, we next investigated whether PTPRU may have altered substrate specificity. Notably, the PTPRQ D1 domain has the same non-canonical aspartate to glutamate substitution in the WPD-loop observed in PTPRU (**SI Appendix, Fig. S1**), and has been previously reported to be a phosphoinositide phosphatase (19). However, PTPRU-D1 exhibited no measurable phosphatase activity against either phosphatidylinositol (PI)-4-phosphate or PI-4,5-bisphosphate (**SI Appendix, Fig. S2A**). While our investigation of protein phosphorylation was primarily focused on residues modified with an O-linked, phosphoester bond (pTyr, pSer, pThr), a recent mass spectrometry study has reported residues modified by an N-linked, phosphoramidate bond (pHis, pLys, pArg, pAsp) as having greater abundance in the cell than pTyr (31). Indeed, human histidine phosphatases have been identified previously (32–34). To test the ability of PTPRU to catalyze hydrolysis of phosphoramidate bonds we used the generic histidine phosphatase substrate imidodiphosphate (PNP; **SI Appendix, Fig. S2B**) (35) but were again unable to detect any activity (**SI Appendix, Fig. S2C**). Together these data support that PTPRU does not catalyze the hydrolysis of phosphoester or phosphoramidate-based substrates.

In the absence of an identifiable substrate, we next sought to determine the molecular mechanism of PTPRU-D1 inactivity. Substitution of the WPD-loop aspartate for glutamate, as seen in PTPRU, is common amongst the D2 pseudophosphatase domains of RPTPs (**SI Appendix, Fig. S3A**) (7). Similarly, mutation of the corresponding Asp (D181) to Glu in the active phosphatase PTP1B results in reduced activity by several orders of magnitude, although it does retain some activity compared with a D181A mutation (36). Mutation of the WPD loop in PTPRK-D1 (D1057E) results in a loss of activity similar to that seen previously for PTP1B, having low residual activity with a *k*_cat_ = 0.0389 s^-1^, corresponding to a ∼115-fold reduction in enzymatic turnover versus WT (**Fig. 1F**). Critically, PTPRU-D1.E1053D, where the Glu has been reverted to the canonical Asp, remained inactive against cellular pTyr (**Fig. 1G**). These data therefore suggest that an Asp to Glu substitution in the WPD loop alone is insufficient to account for complete loss of PTPRU-D1 enzymatic activity.

### Structure of PTPRU-D1

In order to better understand the mechanisms underlying the lack of catalytic activity of PTPRU we determined the structure of the D1 domain. The X-ray crystal structure of PTPRU-D1 was solved by molecular replacement using the PTPRK-D1 domain (37) (PDB ID: 2C7S) and refined to 1.72 Å resolution (Table 1). The overall fold of PTPRU-D1 closely resembles related phosphatase domains (RMSD of 1.014 Å^2^ over 237 Cα atoms with PTPRK, **Fig. 2A**). However, the PTPRU structure reveals several key differences that likely contribute to its catalytic inactivity. The most striking difference is the absence of a folded pTyr recognition loop (**Fig. 2A**). The structure of this loop is well conserved across the PTPs (**Fig. 2B**). In the PTPRU crystal structure residues 904-925 encompassing the pTyr recognition loop region are disordered, resulting in almost complete loss of the pocket that would normally bind the pTyr substrate (**SI Appendix, Fig. S3B**). Consistent with this disorder PTPRU-D1 has reduced protein stability compared to PTPRK-D1, as shown by increased susceptibility to proteolysis (**Fig. 2C**) and a lower global melting temperature (PTPRU-D1 = 48.2°C, PTPRK-D1 = 51.5°C; **Fig. 2D**).

**Fig. 2.**
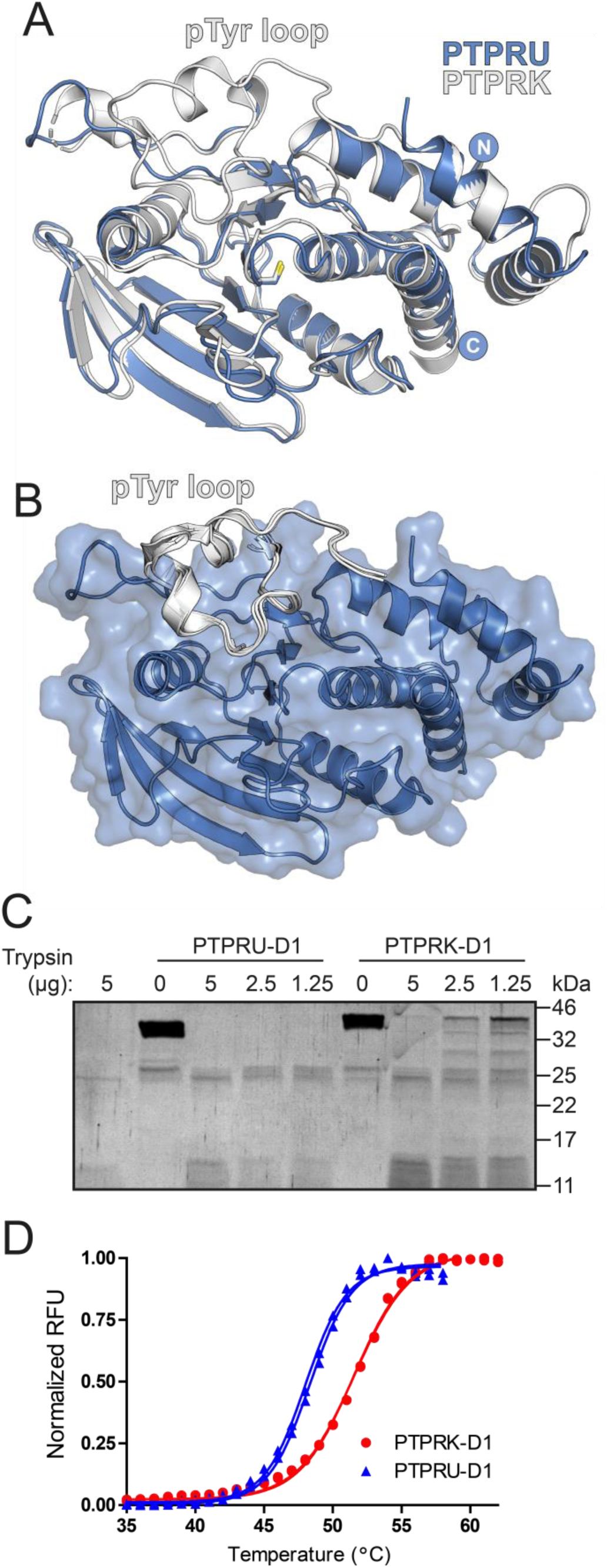
The structure of PTPRU-D1 reveals pTyr recognition loop disorder. *(A)* Ribbon diagram showing the overall structure of the PTPRU-D1 domain (blue) overlaid with PTPRK-D1 (white; PDB ID: 2C7S). *(B)* Transparent surface representation of PTPRU-D1 with superimposed pTyr recognition loops of aligned R2B family D1 domain structures (white ribbons). PDB ID: 2C7S, 2OOQ, 1RPM for PTPRK, PTPRT, PTPRM respectively. *(C)* Limited proteolysis of PTPRU-D1 and PTPRK-D1 with trypsin followed by SDS-PAGE and Coomassie staining. *(D)* Differential scanning fluorimetry thermal profiles of PTPRK-D1 (red) and PTPRU-D1 (blue) recombinant proteins.

**Table 1.** Data Collection and Refinement Statistics. Values in parentheses are for highest-resolution shell.

| **Data collection** | PTPRU-D1 Reduced | PTPRU-D1 Oxidized |
| --- | --- | --- |
| Beamline | I04 | I03 |
| Wavelength (Å) | 0.9795 | 0.9796 |
| Space group | *C* 2 2 2<sub>1</sub> | *C* 2 2 2<sub>1</sub> |
| Cell dimensions |  |  |
| *a,b,c* (Å) | 61.8, 107.9, 88.3 | 62.1, 107.9, 89.3 |
| *a, b, g* (°) | 90, 90, 90 | 90, 90, 90 |
| Resolution (Å) | 53.97 – 1.72 (1.75 – 1.72) | 89.28 – 1.97 (2.00 – 1.97) |
| *R*<sub>merge</sub> | 0.148 (0.904) | 0.128 (1.061) |
| *R*<sub>pim</sub> | 0.043 (0.284) | 0.055 (0.458) |
| CC<sub>1/2</sub> | 0.998 (0.501) | 0.995 (0.527) |
| *I*/*σI* | 9.7 (2.0) | 9.0 (1.6) |
| Completeness (%) | 100 (99.0) | 100 (100) |
| Multiplicity | 12.9 (11.0) | 6.4 (6.3) |
| **Refinement** |  |  |
| Resolution (Å) | 53.97 – 1.72 (1.78 – 1.72) | 46.19 – 1.97 (2.07 – 1.97) |
| No. reflections | 31740 | 21577 |
| *R*<sub>work</sub>/*R*<sub>free</sub> | 0.182/0.201 | 0.214/0.245 |
| No. atoms |  |  |
| Protein | 2020 | 1953 |
| Water | 117 | 94 |
| Other | 2 | 0 |
| *B*-factors |  |  |
| Protein | 29.12 | 37.07 |
| Water | 33.87 | 43.17 |
| Other | 31.41 | - |
| Ramachandran |  |  |
| Favoured (%) | 96.4 | 98.3 |
| Outliers (%) | 0.0 | 0.0 |
| r.m.s. deviations |  |  |
| Bond lengths (Å) | 0.012 | 0.005 |
| Bond angles (°) | 1.110 | 0.863 |
| **PDB ID** | 6SUB | 6SUC |

Another striking difference observed in our PTPRU-D1 structure was the conformation of the PTP loop containing the catalytic cysteine (**Fig. 3A**). The conformation of this loop is highly conserved in classical PTPs (**Fig. 3Ai**), but in PTPRU-D1 this loop is arranged such that the sidechain of T1089 is in close proximity to the active site cysteine (3.0 Å between T1089 OG1 and C1085 SG atoms, **Fig. 3Aii**). Although the majority of the surrounding active site residues are maintained in the canonical conformation, this new loop orientation blocks the catalytic cysteine and would directly interfere with pTyr binding (**Fig. 3Aiii**). A threonine at this position in the PTP loop is unique to the PTPRU-D1 domain (**SI Appendix, Fig. S1**). To probe the importance of this residue for catalysis we created a PTPRU-D1 T1089A mutant and the reciprocal PTPRK-D1 A1093T and tested these in dephosphorylation assays. Introduction of a threonine to the active site was not sufficient to inactivate PTPRK-D1 (**Fig. 3B**). Further, removal of the active site threonine was not sufficient to reactivate PTPRU-D1 and a double mutant E1053D and T1089A also remained inactive (**Fig. 3B**). The combined effect of a disordered pTyr recognition loop and reorientation of the catalytic PTP loop is the loss of key structural features normally required for binding and processing of phosphorylated substrates (**Fig. 3C**).

**Fig. 3.**
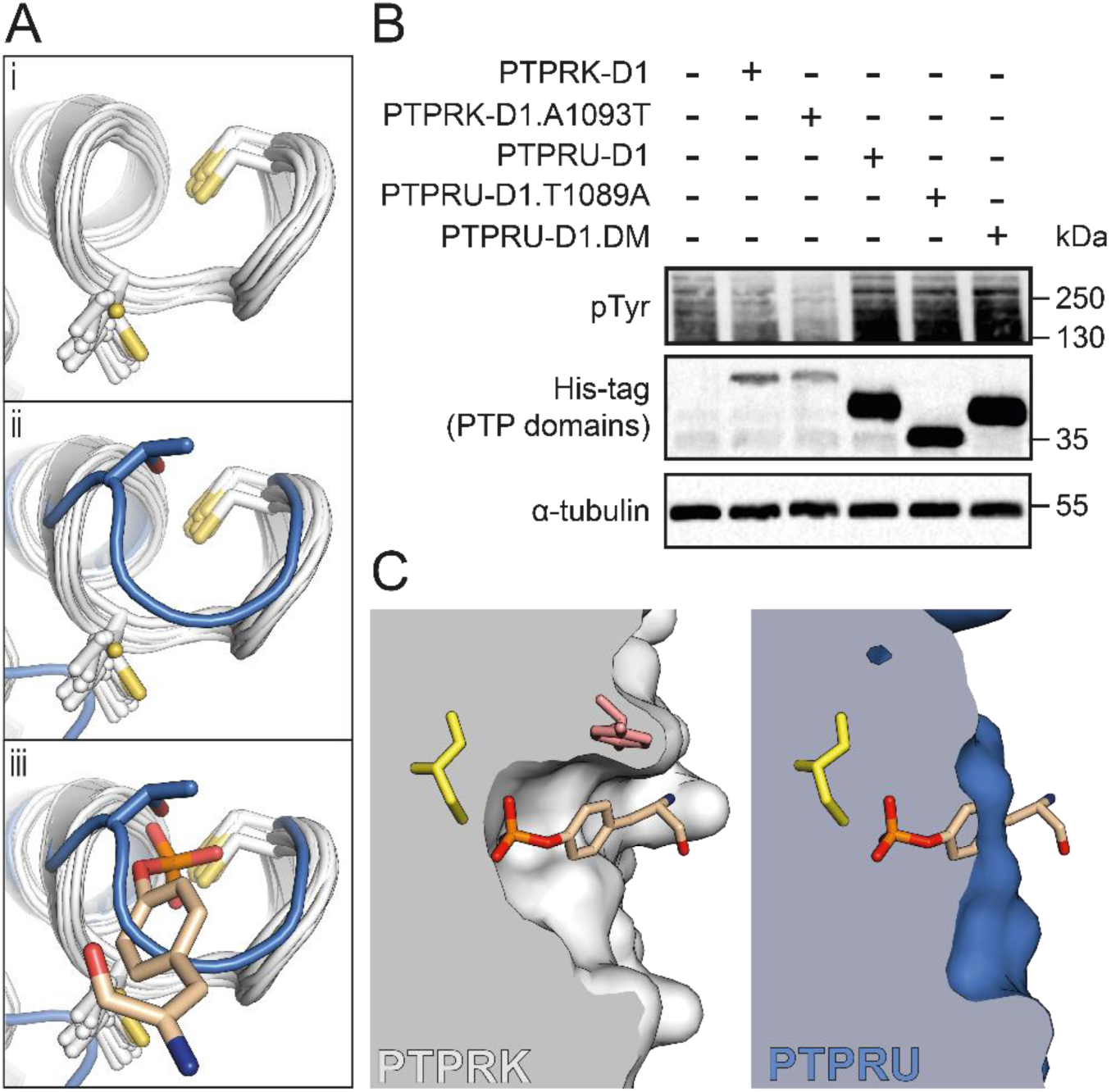
The unique T1089 of PTPRU-D1 blocks the active site pocket. *(A) (i)* Ribbon diagram showing structural alignment of the PTP loop region for all PTP domains with a solved structure. Details of PTPs and corresponding PDB IDs are outlined in **SI Appendix,** Table S1. *(ii)* Overlay of the PTPRU-D1 PTP-loop (blue), which exhibits a distinct conformation. *(iii)* Illustration of pTyr (stick representation) binding in the active site based on structural alignment with PTPN3-Eps15 complex (PDB ID: 4RH5). *(B)* Immunoblot analysis of pervanadate-treated MCF10A lysates incubated with either 0.1 μM PTPRK-D1, PTPRK-D1.A1093T or 5 μM PTPRU-D1, PTPRU-D1.T1089A, PTPRU-D1.E1053D.T1089A (double mutant; DM). NB: All PTP domains are tagged with a 6xHis.TEV.Avi-tag (See Materials & Methods) apart from PTPRU-D1.T1089A which has a 6xHis-tag only, resulting in lower apparent M_W_. *(C) Left,* cross-section through the PTPRK-D1 active site (white surface). Structural alignment with PTPN3-Eps15 complex illustrates how the pTyr substrate would bind within the PTPRK-D1 catalytic pocket, interacting with the catalytic C1089 (yellow) of the PTP loop and the Y917 (pink) of the pTyr recognition loop. *Right,* the equivalent representation of the PTPRU-D1 (blue) demonstrates the structural changes that block substrate binding to the active site.

In PTP1B a secondary pTyr binding site was identified by binding of a peptide containing tandem pTyr residues (38) (PDB ID: 1G1H). This secondary site is proximal to the active site and pTyr binding involves two arginine residues (R24 and R254 in PTP1B). One of the arginine residues, R254, is completely conserved in all D1 sequences except PTPRU where it is uniquely a cysteine (C1121) (7). The loop containing this cysteine in PTPRU-D1 (C1121 to M1127) is poorly ordered and the difficulty of building a single, reliable conformation in the electron density suggests it may adopt multiple conformations. The importance of a cysteine residue in this secondary binding site, in a position resembling that of an active site cysteine, remains unclear.

### Structure of oxidized PTPRU-D1

In an attempt to determine if substrate binding might induce folding of the pTyr recognition loop or rearrangement of the catalytic PTP loop, we soaked PTPRU-D1 crystals with several potential ligands including PO_4_, pTyr and PNP. In none of the datasets collected for these crystal soaks was there any evidence of ligand binding in the active site or any induced folding of the pTyr recognition loop. However, these crystals, collected 4 weeks after the initial datasets, had clearly undergone oxidation resulting in the formation of a disulfide bridge between the catalytic C1085 and the vicinal “backdoor” cysteine, C998 (**Fig. 4Ai**). This alternate conformation involving disulfide bond formation with nearby cysteines has been observed for several other related phosphatases (**Fig. 4Aii**) (39–42). This disulfide formation upon oxidation has been proposed to protect the catalytic cysteine from oxidative damage and/or function as a redox-sensitive mechanism for reversible PTP inactivation (39–42). The formation of this disulfide in PTPRU-D1 destabilizes the conformation of adjacent residues in the PTP-loop as there is no clear density in which to model S1086-G1088 (**Fig. 4B**). The nearby loop (C1121-M1127) previously implicated as a secondary pTyr binding site is also further destabilized in this oxidized structure as evidenced by a lack of electron density for this region. Non-reducing SDS-PAGE of PTPRU-D1 recombinant protein in the presence of hydrogen peroxide results in a mobility shift consistent with disulfide formation in solution, which is completely reversed under reducing conditions (**Fig. 4C**). While PTPRK-D1 conserves this “backdoor” cysteine, it does not undergo detectable disulfide formation under the same conditions (**Fig. 4C**). Thus, the catalytic cysteine of PTPRU-D1 can undergo reversible oxidation, involving intramolecular disulfide formation, identifying this domain as a redox-sensitive pseudophosphatase.

**Fig. 4.**
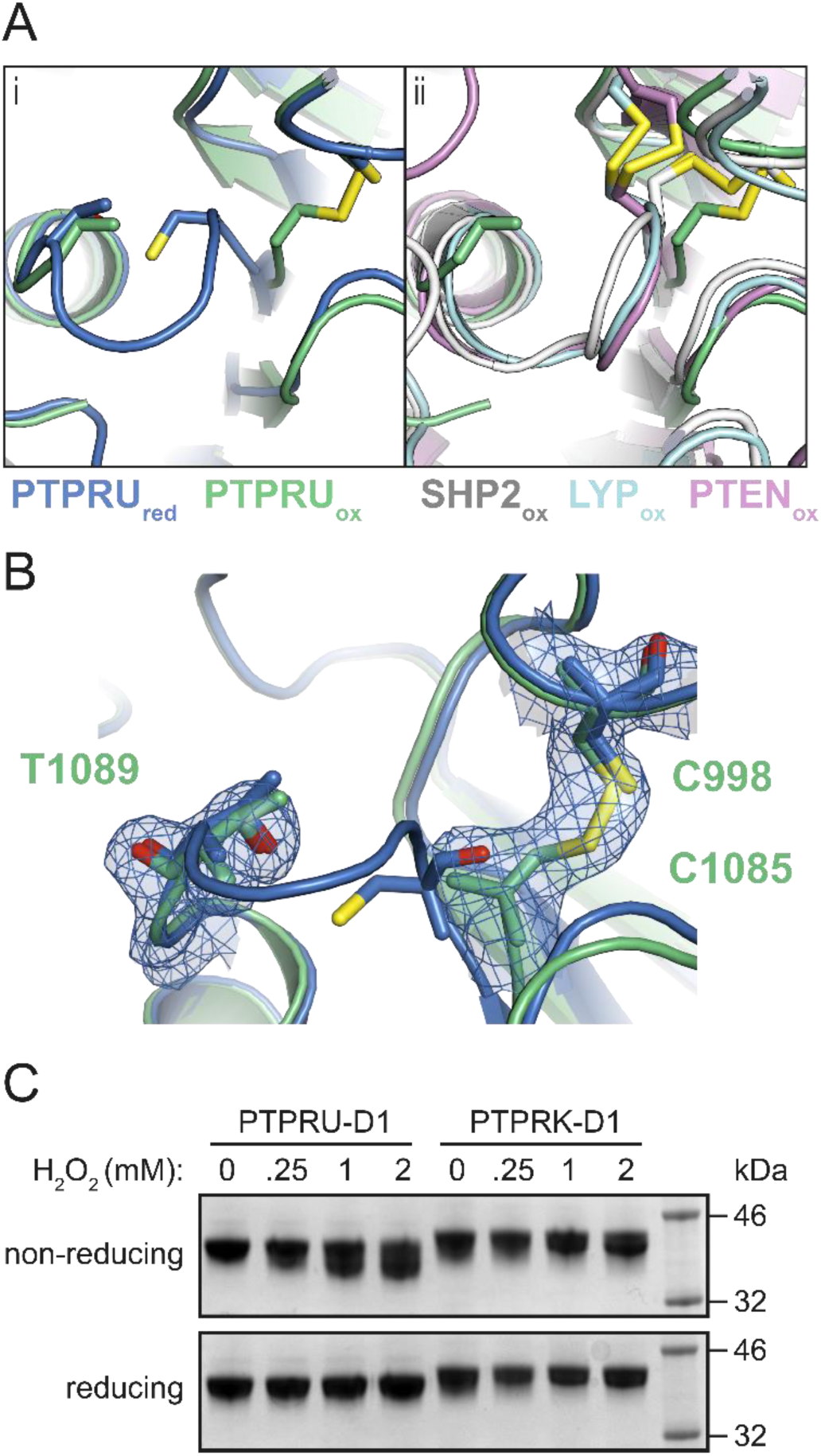
The PTPRU D1 domain forms an intramolecular “backdoor” disulfide bond. *(A) (i)* Ribbon diagram of the reduced PTPRU-D1 PTP-loop region (PTPRU_red_, blue) overlaid with PTPRU-D1 oxidized (PTPRU_ox_, green). *(ii)* Ribbon diagram of the PTPRU_ox_ structure (green) overlaid with previously solved “backdoor” disulfide containing structures (SHP2, white; LYP, cyan; PTEN, violet; PDB IDs 6ATD, 3H2X and 5BUG respectively). *(B)* Reduced and oxidized models of PTPRU-D1 colored as in *(A)(i)*, showing electron density (2*F*_o_-*F*_c_ contoured at 0.8 e^-^/Å^3^, blue) for the “backdoor” disulfide and PTP-loop of the oxidized form. *(C) In vitro* oxidation of PTPRU and PTPRK D1 domains by hydrogen peroxide followed by reducing and non-reducing SDS-PAGE and Coomassie staining.

### PTPRU interacts with PTPRK substrates

To understand the role of PTPRU in signaling, we exploited our previous observation based on domain-swapping intracellular domain (ICD) chimeras showing that the PTPRK, but not PTPRM, D2 domain was critical for recognition of Afadin (3), a reported PTPRU interactor (29). Although we were unable to purify full PTPRU-ICD (D1+D2), we were able to generate an *in vivo* biotinylated chimeric ICD consisting of the active PTPRK-D1 and the PTPRU-D2 domain (**Fig. 5A; SI Appendix, Fig. S4A and S4B**). We then tested the ability of chimeric proteins to bind and dephosphorylate PTPRK substrates. To probe protein-substrate interactions, we conjugated biotinylated chimeric proteins to streptavidin beads for *in vitro* pull-downs from pervanadate-treated cell lysates followed by immunoblotting. While the PTPRK and PTPRM substrate p120-Catenin (p120^Cat^) (3) could interact with all chimeras regardless of D2 domain, we found that unlike PTPRM-D2, the PTPRU-D2 domain is sufficient for binding to Afadin (**Fig. 5B**). Consistent with this interaction data, immunoprecipitation of tyrosine phosphorylated proteins from dephosphorylation assays confirmed the PTPRU-D2 domain as being sufficient to recruit Afadin for dephosphorylation by the active PTPRK-D1 domain (dephosphorylated proteins are depleted in pTyr IPs and/or enriched in supernatants; **Fig. 5C**). Our data suggest that in cells PTPRU will bind but not dephosphorylate PTPRK substrates.

**Fig. 5.**
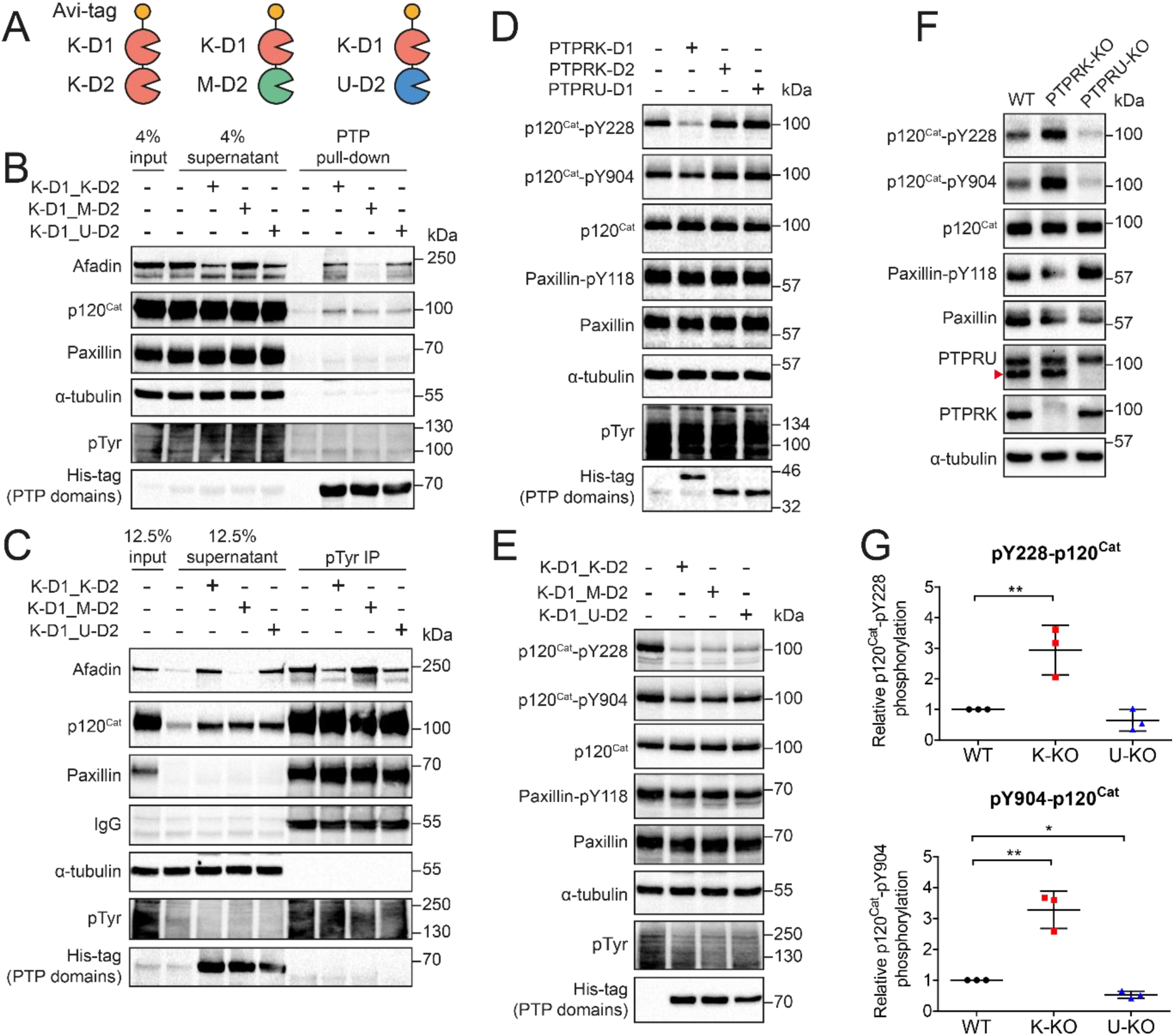
PTPRU binds but does not dephosphorylate PTPRK cellular substrates. *(A)* Schematic diagram of chimeric tandem PTP domains used in this study. *(B)* Immunoblot analysis of recombinant protein pull-downs from pervanadate-treated MCF10A lysates, using the indicated chimeric tandem PTP domains. *(C)* Immunoblot analysis of pTyr immunoprecipitations (IPs) from dephosphorylation assays. Pervanadate treated MCF10A lysates incubated for 1.5 hours at 4°C with 300 nM of the indicated chimeric tandem PTP domains. Dephosphorylated proteins are depleted from IPs and enriched in supernatants. *(D)* Immunoblot analysis of pervanadate-treated MCF10A lysates incubated for 1.5 hours at 4°C with 100 nM of PTPRK-D1, PTPRK-D2 or PTPRU-D1 recombinant domains. *(E)* Immunoblot analysis of pervanadate-treated MCF10A lysates incubated for 1.5 hours at 4°C with 300 nM of the indicated chimeric tandem PTP domains. *(F)* Immunoblot analysis of confluent WT, PTPRK-KO and PTPRU-KO MCF10A cell lysates, harvested 24 hours post media change. *(G)* Densitometry of p120^Cat^ phosphorylation levels from *(F)*, normalized against total p120^Cat^ protein levels. Error bars represent ± SEM of *n* = 3. Un-paired student’s t-test: p < 0.05 (*), p < 0.005 (**).

Previously we have identified specific p120^Cat^ pTyr residues (Y228, Y904) which are dephosphorylated by PTPRK and PTPRM D1 domains, and are hyperphosphorylated in PTPRK-KO cells (3). We confirmed by in lysate dephosphorylation assays that while all PTPRK-D1 containing chimeras show dephosphorylation of pY228 and pY904, PTPRU-D1 cannot dephosphorylate these p120^Cat^ sites (**Fig. 5D and 5E**). To investigate the cellular consequence of PTPRU binding to PTPRK substrates, we generated CRISPR-Cas9 mediated PTPRU-KO MCF10A cells. As expected, we were able to observe hyperphosphorylation of p120^Cat^-pY228 and -pY904 in PTPRK-KO cells vs WT (**Fig. 5F**). Strikingly, deleting PTPRU resulted in hypophosphorylation of both p120^Cat^-pY228 and -pY904 vs WT levels (**Fig. 5F** and **Fig. 5G**). Taking our interaction and dephosphorylation data together, this supports a mechanism in which PTPRU can bind substrates and protect them from dephosphorylation by related phosphatases.

## DISCUSSION

The receptor PTPRU exhibits divergent sequences in key catalytic motifs including a highly divergent pTyr recognition loop sequence, a unique Thr within the PTP loop and a Glu substitution in the WPD loop. Here we show that PTPRU does not possess detectable phosphatase activity against a range of substrates or in dephosphorylation assays with cell lysates. Our structural data identify multiple features that would disrupt both substrate binding and catalytic activity. Despite its inactivity, we demonstrate that PTPRU can bind to key proteins previously reported as substrates for its catalytically active paralog PTPRK. This supports a role for PTPRU as a scaffold that competes with active phosphatases at the plasma membrane to locally influence tyrosine phosphorylation dynamics and potentially cell-cell adhesion.

Previous studies have shown that WPD to WPE mutations in several active phosphatases results in significant reduction in enzyme activity (13, 36). We show a similar substantial decrease in PTPRK activity following the introduction of the WPE sequence change. However, mutation of the PTPRU WPE loop to the canonical sequence was not sufficient to rescue any detectable phosphatase activity. Our structure of the PTPRU-D1 reveals that there are two key structural changes within this domain that alter the substrate binding pocket and therefore are likely to explain the lack of phosphatase activity. The first is the disordered pTyr recognition loop. The absence of electron density for this loop was unexpected as, although the PTPRU sequence is highly divergent, it does retain a conserved arginine (R918) that in related structures binds back into the main PTP fold and interacts with residues that are conserved in PTPRU-D1. Not only does loss of this ordered loop drastically alter the shape of the substrate binding pocket, it also contributes to decreased protein stability as demonstrated by the increased susceptibility of PTPRU to proteolysis and its lower melting temperature relative to PTPRK-D1. The second key structural change in PTPRU-D1 relates to the catalytic PTP loop, which has undergone a substantial rearrangement resulting in the occlusion of the substrate binding site, blocking the catalytic cysteine. Interestingly, the introduction of a Thr into the equivalent position of PTPRK-D1 does not reduce the phosphatase activity of this domain and mutation of the PTPRU Thr to Ala does not induce detectable PTPRU activity. Furthermore, the introduction of two mutations, T1089A and E1053D, into the PTPRU-D1 PTP and WPD loops, respectively, were still unable to rescue any activity. This combination of biochemical and structural analysis demonstrates that there are multiple mechanisms contributing to the absence of phosphatase activity in PTPRU-D1.

An intriguing feature of the inactive D2 pseudophosphatase domains in the RPTP family is the retention of the catalytic cysteine residue. The conservation of this residue in inactive domains raises the question of whether it plays an alternative, non-catalytic role in these domains. Our structure of oxidized PTPRU-D1 demonstrates that this cysteine has the capacity to form a “backdoor” disulfide similar to that seen in SHP2, LYP and PTEN phosphatases. In these enzymes, the formation of a disulfide bond has been attributed to the need to protect the catalytic cysteine from irreversible oxidative damage or to allow reversible redox-sensitive inactivation. Our observation here of a similar intramolecular disulfide in an inactive pseudophosphatase domain suggests that the proposed roles for the disulfide formation may extend beyond the modulation of enzyme activity. Indeed, previous studies on the PTPRA D2 domain suggest oxidation can promote an intermolecular disulfide bond (17), or a conformational change that is translated to the extracellular domain (43).

In addition to promoting a “backdoor” disulfide, PTP oxidation can induce chemical modification of the catalytic cysteine. One such modification is the formation of a sulfenylamide intermediate, as demonstrated for PTP1B (44, 45). This modification involves the sidechain of the catalytic cysteine forming a covalent link to the backbone nitrogen of an adjacent residue, resulting in a substantial change to the conformation of the catalytic PTP loop. Interestingly, this loop conformation is highly similar to that seen in PTPRU-D1. In PTP1B this renders the enzyme inactive but is reversible upon reduction and is proposed to be a protective intermediate during redox-regulated inhibition. This conformation of the PTPRU-D1 PTP loop is not induced by oxidation, it is instead present in the reduced form rendering the enzyme unable to bind phosphotyrosine. This suggests PTPRU has evolved to adopt an inactive conformation, even under reducing conditions.

The ICDs of other members of the R2B family comprise an active membrane-proximal D1 domain and an inactive membrane-distal D2 domain. Despite the sequence divergence of PTPRU-D1 from its paralogs (**SI Appendix, Fig. S1**) and its lack of catalytic activity, this domain still possesses higher sequence identity to D1 domains than to D2 domains (69% sequence identity with R2B family D1 domains, 28% identity with R2B family D2 domains; **SI Appendix, Table S2**). Therefore, rather than PTPRU possessing two inactive D2 domains, it retains a *bona-fide* D1 and D2 domain topology similar to that of related enzymes but with a D1 domain that has diverged to be catalytically inactive. By using chimeric ICDs containing D1 and D2 domains from PTPRU, PTPRK and PTPRM in cell-based dephosphorylation assays we show that the D2 domain of PTPRU can recruit substrates for dephosphorylation by the active D1 domain of PTPRK. This ability to bind substrates that overlap with active phosphatases, combined with the lack of phosphatase activity of PTPRU suggests that the likely role of PTPRU in cells is to act as a decoy receptor that sequesters substrates protecting them from dephosphorylation. In support of this, we find that genetic deletion of PTPRU leads to a reduction in phosphorylation levels of p120^Cat^. In this way, PTPRU may modify cell signaling by altering the rate or extent of tyrosine dephosphorylation by related, active RPTP family members. The absence of enzyme activity demonstrated here for PTPRU does not diminish its importance but highlights a new pseudophosphatase function in cell signaling.

## MATERIALS AND METHODS

### Protein Expression and Purification

Constructs for expressing PTPRU, PTPRK D1 and D2 domains and chimeric intracellular domains (ICDs) were cloned with an N-terminal 6xHis-tag alone or with a tandem N-terminal 6xHis and Avi-tags. Point mutations were introduced using site-directed mutagenesis and sequence verified. All proteins were expressed in *E. coli* BL21 (DE3) Rosetta cells and purified using nickel-affinity and size exclusion chromatography.

### Crystallization, Data Collection and Structure Determination

For crystallization trials, PTPRU-D1 protein was further purified by anion exchange chromatography and concentrated to 9.6 mg/ml in 50 mM Tris-HCl, pH 8, 300 mM NaCl, 5% glycerol, 5 mM DTT. Crystals were grown by sitting-drop vapor diffusion against a reservoir of 0.1 M Bicine, pH 9, 1 M lithium chloride, 20% (w/v) polyethylene glycol 6000. Crystals were cryoprotected in reservoir solution supplemented with 20% (v/v) glycerol and flash-cooled by plunging into liquid nitrogen. Diffraction data were collected at the Diamond Light Source (DLS) beamlines I03 and I04 and processed using the automated data processing pipelines at DLS (46) (**Table 1**). The structure of PTPRU-D1 was solved by molecular replacement with PTPRK-D1 (PDB ID: 2C7S), model building was carried out using COOT (47) and refined using phenix.refine (**Table 1**) (48).

Detailed Materials and Methods for all experiments are provided in **SI Materials and Methods**.

## Data availability

The atomic coordinates and structure factors have been deposited in the Protein Data Bank, www.pdb.org under accession codes 6SUB (PTPRU-D1 reduced form) and 6SUC (PTPRU-D1 oxidized form). Other data and reagents are available from the corresponding authors upon reasonable request.

## ACKNOWLEDGEMENTS

We acknowledge Diamond Light Source for time on beamlines I03 and I04 under proposal MX15916. Remote access was supported in part by the EU FP7 infrastructure grant BIOSTRUCT-X (Contract No. 283570). We thank Stephen Graham for assistance with the crystallography work and the CIMR flow cytometry core facility for their assistance in cell-sorting. IMH is supported by a CIMR PhD studentship. M.K. acknowledges funding from the German Research Foundation (DFG) under Germany’s Excellence Strategy (CIBSS – EXC-2189 – Project ID 390939984 and BIOSS – EXC-294). HJS and GWF are funded by a Sir Henry Dale Fellowship jointly funded by the Wellcome Trust and the Royal Society (109407). JED is funded by a Royal Society University Research Fellowship (UF150682).

## SUPPLEMENTARY INFORMATION

### MATERIALS AND METHODS

#### Plasmids and constructs

Amino acid (aa) numbering used throughout is based on the following sequences; PTPRU; UniProt ID: Q92729-1; PTPRK; UniProt ID: Q15262-3; PTPRM; UniProt ID: P28827-1; PTPRT; UniProt ID: O14522-3. For bacterial expression, human cDNA sequences encoding PTPRU-D1 (aa. 871-1153), PTPRU-D2 (aa 1150-1446), PTPRK-D1 (aa 865-1157), PTPRK-D2 (aa 1154-1446) were subcloned into a modified pET-15b expression vector in frame with an N-terminal hexahistidine-tag followed by a TEV protease recognition site and Avi-tag (1) (His6.TEV.Avi-tag). Chimeric ICDs were generated by subcloning of PTPRK-D1(aa 865-1153).BstBI.PTPRK-D2(aa 1154-1146) and PTPRK-D1(aa 865-1153).BstBI.PTPRU-D2(aa 1150-1446) in to pET-15b.His6.TEV.AviTag. For crystallization studies PTPRU-D1 was subcloned into a modified pET-15b expression vector in frame with an N-terminal His6 tag only. Mutations were introduced by site-directed mutagenesis using polymerase chain reaction with Phusion Hot Start II DNA polymerase.

#### Antibodies

The antibodies used for immunoblot analysis in this study are as follows. All antibodies were used at a dilution of 1:1000 in TBS-T (20 mM Tris-HCl, pH 7.6, 137 mM NaCl, 0.2% [v/v] Tween-20) with 3% (w/v) BSA unless otherwise indicated. Rabbit anti-pTyr (Cat#8954), mouse anti-HisTag (Cat#2366), rabbit anti-Paxillin (Cat#12065), rabbit anti-phospho-Paxillin (Y118; Cat#2541), rabbit anti-phospho-p120-Catenin (Y228; Cat#2911), and rabbit anti-phospho-p120-Catenin (Y904; Cat#2910) primary antibodies were all purchased from Cell Signalling Technology. Mouse anti-Afadin (Cat#610732) and mouse anti-p120-Catenin (Cat#610133) primary antibodies were purchased from BD Transduction Laboratories. Mouse anti-PTPRM (PTPRU cross-reactive; Cat#sc-56959) (2) primary antibody was purchased from Santa Cruz. Rabbit anti-PTPRK primary antibody was generated in a previous study (1). Mouse anti-alpha-tubulin primary antibody was purchased from Sigma Aldrich (Cat#T6199). HRP conjugated anti-mouse (Cat#711-035-151) and anti-rabbit (Cat#711-035-152) secondary antibodies (1:5000 in TBS-T) were purchased from Jackson Immuno-Research.

#### Protein expression and purification

*Escherichia coli* BL21(DE3) Rosetta cells transformed with the relevant expression construct were cultured in 2X TY medium at 30°C until OD_600_ = 0.6. Routinely, 1-2 mg of recombinant PTP was obtained per 1 L culture. Protein expression was induced with 1 mM isopropyl-thio-β-D-galactopyranoside for 18 hours at 20°C. For biotinylated Avi-tag constructs, 200 μM D-biotin (Sigma Aldrich) was added at the point of induction. After a freeze-thaw cycle, bacterial pellets were resuspended in ice-cold lysis buffer (50 mM Tris, pH 7.4 [PTPRK domains]/pH 8 [PTPRU domains], 500 mM NaCl, 5% glycerol, 0.5 mM TCEP) and lysed using high-pressure cell disruption (Constant Systems Ltd). Cell lysates were clarified by centrifugation at 40,000 xg, 30 min. Cleared lysates were incubated with Ni-NTA agarose beads (Qiagen) for 1 h at 4°C. Ni-NTA beads were packed in to a 10 ml gravity-flow column and equilibrated with 10 bed volumes of purification buffer (for PTPRU constructs; 50 mM Tris-HCl, pH 8, 500 mM NaCl, 5% glycerol, 5 mM DTT, for PTPRK constructs; 50 mM Tris-HCl, pH 7.4, 500 mM NaCl, 5% glycerol, 5 mM DTT) containing 5 mM imidazole. Ni-NTA beads were then washed with 20 bed volumes of purification buffer containing 20 mM imidazole, followed by elution in purification buffer containing 250 mM imidazole. Eluted protein was further purified by size-exclusion chromatography on a HiLoad Superdex 200 pg 16/600 column (GE Healthcare) equilibrated in purification buffer. For crystallization-quality PTPRU-D1 domain, protein was buffer exchanged by iterative concentration/dilution in a 10K MWCO centrifugal filter unit (Merck Millipore) against low-salt buffer (50 mM Tris-HCl, 10 mM NaCl, pH 8, 5% glycerol, 5 mM DTT) until a final NaCl concentration <15 mM. Protein was further purified by anion exchange chromatography on a MonoQ 5/50 GL column (GE Healthcare) equilibrated in low-salt buffer and bound protein was eluted by a linear 20 ml gradient against high-salt buffer (50 mM Tris-HCl, pH 8, 1M NaCl, 5% glycerol, 5 mM DTT). Protein purity was assessed by SDS-PAGE and staining with Coomassie (Instant Blue, Expedeon).

#### Crystallization

Crystallization experiments were performed in 96-well nanolitre-scale sitting drops (200 nl of 9.6 mg/ml PTPRU-D1 plus 200 nl of precipitant) equilibrated at 20°C against 80 μl reservoirs of precipitant. Diffraction quality crystals grew against a reservoir of 0.1 M Bicine, pH 9, 1 M lithium chloride, 20% (w/v) polyethylene glycol 6000. For the oxidised structure, PTPRU-D1 crystals were soaked in 1 mM pTyr (Sigma Aldrich) for 3 min. Crystals were cryoprotected in reservoir solution supplemented with 20% (v/v) glycerol and flash-cooled by plunging into liquid nitrogen.

#### X-ray data collection and structure solution

X-ray diffraction data were recorded at Diamond Light Source (DLS) beamlines I03 and I04. Datasets were collected at λ = 0.9795 Å. Diffraction datasets were indexed and integrated using the automated data processing pipeline available at DLS, implementing XIA2 DIALS for the reduced dataset and XIA2 3dii for the oxidized dataset (3) then scaled and merged using AIMLESS (4). Resolution cut-off was determined by CC_1/2_ > 0.5 and I/σl > 1.5. The initial structure was solved by molecular replacement using Phaser (5), with human PTPRK-D1 (6) (PDB ID: 2C7S) as a search model. Further refinements were performed using COOT (7) and phenix.refine (8). Graphical figures of the PTPRU-D1 structure were rendered in PyMOL (Schrödinger, LLC).

#### pNPP phosphatase activity assay

Recombinant PTP domains were made up to 50 μL volumes in a 96-well microplate format in PTP buffer (50 mM Tris-HCl, pH 7.4, 150 mM NaCl, 5% glycerol, 5 mM DTT, 100 μg/ml BSA). Serial dilutions of pNPP substrate (0-40 mM, New England Biolabs) were performed in PTP buffer. Reaction plates containing both enzyme and substrate dilutions were incubated at 30°C for 30 min prior to addition and mixing of 50 μL pNPP substrate to initiate reactions. Product formation was monitored for 15 min at 30°C by measuring absorption at 405 nm in a Spectramax M5 plate reader (Molecular Devices), followed by fitting to a 4-nitrophenyl (Sigma) standard curve of known concentration. Data were fitted using linear regression in GraphPad Prism to determine initial enzymatic rates (V_0_). V_0_ values at various substrate concentrations were fitted using non-linear regression and kinetic parameters (V_max_ and K_m_) calculated from the Michaelis-Menten equation in GraphPad Prism. kcat values were calculated using the equation k_cat_ = *V*_max_ / [E_T_].

#### BIOMOL Green phosphatase assay

Recombinant PTP domains were made up to 20 μl volumes in a 96-well microplate format in reaction buffer (50 mM Tris-HCl, pH 7.4, 150 mM NaCl, 5% glycerol, 5 mM DTT). 30 μl of 100 μM pTyr peptides (DADE-pY-LIPQQG and END-pY-INASL) or imidodiphosphate was added to protein wells to initiate reactions. Reactions were allowed to proceed for 15 min before termination by addition of 100 μl BIOMOL Green reagent (Enzo). Liberated phosphate was measured by absorbance at 360 nm in a Spectramax M5 plate reader (Molecular Devices), followed by interpolation to a standard curve of known phosphate concentration. Serial dilutions of phosphate were performed in reaction buffer using 800 μM phosphate standard (Enzo).

#### Phosphoinositide phosphatase activity assay

The commercially available Enzchek phosphate assay kit (Thermo Fisher Scientific) was used to measure the release of phosphate from the phosphoinositides lipids PI(4,5)P2 and PI(4)P. The kit was used according to the manufacturer’s instructions. Briefly, PTPRU-D1 (3 µM) or the positive control PRL-3 (6 µM) were incubated in Enzchek reaction buffer containing 50 mm Tris-HCl pH 7.5, 2 mM MgCl2 plus 500 mM NaCl (only for PTPRU-D1) and 5 mM DTT. The reaction was initiated upon addition of each lipid substrate at a final concentration of 100 µM. The assay was conducted in triplicates at 37° C in a BioTek Synergy H1 plate reader for 45 minutes and the release of phosphate was monitored measuring the absorbance at 360 nm over time. For every phosphoinositide substrate, a control without enzyme for blank subtraction was also measured. The data are represented as mean +/− SD.

#### Cells and cell culture

MCF10A cells were obtained from the American Type Culture Collection (ATCC). MCF10A cells were cultured in a previously described MCF10A growth medium (9) consisting of 1:1 DMEM:Ham’s F12 supplemented with 5% (v/v) horse serum, 0.5 mg/ml hydrocortisone, 20 ng/ml EGF, 100 ng/ml cholera toxin and 10 mg/ml insulin. Cells were cultured at 37°C in a humidified 5% CO_2_ atmosphere in 75 cm^2^ vented flasks.

#### Preparation of sodium pervanadate

To create a 50 mM pervanadate stock solution, 30% H_2_O_2_ was first diluted to 0.3% H_2_O_2_ in 20 mM HEPES, pH 7.3. 50 μl dilute H_2_O_2_ added to 490 μl 100 mM sodium orthovanadate (Alfa Aesar) and 450 μl H_2_O, then mixed by gentle inversion and incubated at RT for 5 min. After incubation, excess H_2_O_2_ was quenched by the addition of a small amount of catalase (using 200 μl pipette tip) and mixed by gentle inversion. Pervanadate was freshly prepared and used immediately to avoid decomposition.

#### Generation of pervanadate-treated lysates

3 x 10^6^ MCF10A cells were seeded in 10 cm dishes and cultured to complete confluence (4 days). Media was then aspirated and cells treated with 8 ml of fresh growth medium containing 1 mM sodium pervanadate for 30 min at 37°C, 5% CO_2_. Cells were then placed on ice and washed twice with ice-cold PBS. Cells were lysed in 600 μl per 10 cm dish of ice-cold lysis buffer (50 mM Tris-HCl, pH 7.4, 150 mM NaCl, 10% [v/v] glycerol, 1% [v/v] Triton X-100, 1 mM EDTA, 5 mM iodoacetamide [IAA], 1 mM sodium orthovanadate, 10 mM NaF, 1X EDTA-free protease inhibitor) by orbital shaking on ice, in the dark, at 4°C for 30 min. Lysates were collected by cell-scraping and treated with 10 mM DTT on ice for 15 min. Note: EDTA was used to chelate excess vandate and IAA for alkylation of PTP active site cysteines. Lysates were then cleared by centrifugation at 14000 xg, 15 min at 4°C. Supernatants were transferred to a fresh tube and snap frozen in liquid N_2_ for storage at −80°C until use.

#### In lysate dephosphorylation assays

Recombinant PTP domains (0-5 μM final concentration, as indicated) were mixed with 50 μl (800 μg) of freshly thawed pervanadate treated cell lysate in a final volume of 400 μl made up in ice-cold 150 mM wash buffer with 5 mM DTT (20 mM Tris-HCl, pH 7.4, 150 mM NaCl, 10% [v/v] glycerol, 1% [v/v] Triton X-100, 1 mM EDTA, 5 mM DTT). Reactions were then incubated with rotation for 24 hrs at 4°C unless stated otherwise. For time-course experiments, at each timepoint 72 μl of reaction was removed and added to 1.5 μl 20% (w/v) SDS (0.4% [w/v] SDS final) to stop reactions. 26 μl (∼50 μg) of each sample was then mixed with 6.5 μl 5X sample loading buffer and stored at −20°C prior to SDS-PAGE and immunoblot analysis.

#### SDS-PAGE and immunoblotting

Total protein concentrations of cell-lysates were quantified by bicinchoninic acid assay and 25-50 μg of lysate mixed in an appropriate volume of 5X sample loading buffer (0.25 M Tris-HCl, pH 6.8, 10% [w/v] SDS, 20% [v/v] glycerol, 0.1% [w/v] bromophenol blue, 10% [v/v] β-mercaptoethanol). Samples were then incubated at 92°C for 5 min and resolved by SDS-PAGE on a 10% SDS-polyacrylamide gel. Total protein was transferred to 0.2 μm reinforced nitrocellulose membranes by wet transfer and blocked in 5% (w/v) skimmed-milk in TBS with 0.2% TWEEN-20 (TBS-T; 20 mM Tris-HCl, pH 7.6, 137 mM NaCl, 0.2% [v/v] Tween-20) for 20-60 min. Membranes were then rinsed once with TBS-T before incubation with appropriate primary antibody in 3% (w/v) BSA/TBS-T overnight at 4°C. Membranes were washed 3 x 10 min in TBS-T and then incubated with the appropriate species specific HRP-conjugated anti-IgG secondary antibody for 1 h at RT. Following 3 x 10 min washes in TBS-T, membranes were developed using EZ-ECL solution (Geneflow) and imaged using a ChemiDoc MP imaging system (Bio-Rad). 2D-densitometry for quantification was carried out in FijI (10)

#### Limited proteolysis

All steps were performed on ice unless otherwise stated. 100 μg vials of trypsin protease (Thermo Fisher Scientific) were reconstituted at 1 mg/ml in proteolysis buffer (50 mM Tris-HCl, pH 7.4, 150 mM NaCl, 10% glycerol, 1% Triton X-100, 1 mM EDTA, 5 mM DTT). 4 μg of recombinant protein was incubated with 0, 1.25, 2.5 or 5 μg trypsin in a total volume of 25 μl made up in proteolysis buffer. Reactions were incubated on ice for 30 min and terminated by the addition of 10 μl 5X sample loading buffer. Samples were immediately incubated at 92°C for 5 min and resolved by SDS-PAGE on a 16% SDS-polyacrylamide gel. Protein was visualized by Coomassie staining and gels imaged using a ChemiDoc MP imager.

#### Differential scanning fluorimetry

Differential scanning fluorimetry was performed using Protein Thermal Shift Dye kit (Thermo Fisher Scientific) as per manufacturer’s protocol in a ViiA-7 Real-time PCR system (Thermo Fisher Scientific). Reaction mixes consisting of 2 μg recombinant protein and 1X Protein Thermal Shift Dye were made up to a total volume of 20 μl in purification buffer. Samples were then heated on a 1°C/s gradient from 25-95°C and protein unfolding at each temperature monitored by measurement of fluorescence at 580/623 nm (excitation/emission). Fluorescent signal vs. temperature was fitted to a non-linear Boltzman-sigmoidal regression in Graphpad Prism, with the T_m_ calculated from the inflection point of the fitted curve.

#### Reversible oxidation of recombinant protein

All steps were performed on ice unless otherwise stated. 10 μg of recombinant protein was mixed with either 0, 0.25, 1 or 2 mM H_2_O_2_ in a total reaction volume of 50 μl in ice-cold oxidation buffer (50 mM Tris-HCl, pH 7.4, 150 mM NaCl, 5% glycerol). Reactions were then incubated on ice for 30 min. Each reaction was then split equally (25 μl, 5 μg) and incubated at 95°C for 5 min with 7.5 μl 5X sample loading buffer with (reducing) or without (non-reducing) β-mercaptoethanol. Reduced and non-reduced samples were then resolved by SDS-PAGE on separate NuPage 4-12% bis-tris gels using 1X MOPS running buffer and visualized by staining with Coomassie. Gels imaged using a ChemiDoc MP imager.

#### Assessment of recombinant protein biotinylation

10 μg of *in vivo* biotinylated recombinant protein was boiled for 5 min at 95°C in an appropriate volume of 5X sample loading buffer. Samples were cooled to RT before addition of a 3-fold molar excess of streptavidin and incubated for 5 min at RT. Protein was then resolved by SDS-PAGE on a NuPAGE 4-12% Bis-Tris gel and visualized by staining with Coomassie stain. After destaining with ddH_2_O, gels were imaged on a Bio-Rad ChemiDoc MP imager and total levels of biotinylated protein quantified by 2D-densitometry in FijI (10).

#### Recombinant protein pull downs

50 μg of biotinylated Avi-tag recombinant PTP domains were bound to 167 μl of streptavidin-coated magnetic beads (New England Biolabs) made up to a total volume of 500 μl in ice-cold purification buffer (50 mM Tris, pH 7.4, 150 mM NaCl, 5% [v/v] glycerol, 5 mM DTT) at 4°C for 1.5 h with rotation. Beads were collected using a magnetic stand and washed 3 times in ice-cold purification buffer, followed by two washes with ice-cold 150 mM NaCl wash buffer (20 mM Tris-HCl, pH 7.4, 150 mM NaCl, 10% [v/v] glycerol, 1% [v/v] Triton X-100, 1 mM EDTA). Bead conjugated PTP domains were then blocked in ice-cold 5% (w/v) BSA/150 mM NaCl wash buffer at 4°C for 1 h with rotation. Pervanadate-treated cell lysates were pre-cleared by incubation with streptavidin magnetic beads (0.67 mg beads per ml of lysate) at 4°C for 1 h with rotation. Blocked PTP domains were collected on a magnetic stand and washed twice with ice-cold 150 mM NaCl wash buffer. 1 mg of pre-cleared pervanadate-treated lysate (250 μl) was incubated with PTP domain-bound beads in a total volume of 1 ml 150 mM NaCl wash buffer at 4°C for 1.5 h with rotation. At 4°C, beads were collected on a magnetic stand and supernatants removed. Bead bound protein was then washed twice by resuspension in ice-cold 150 mM NaCl wash buffer, followed by one wash in ice-cold 500 mM NaCl wash buffer (20 mM Tris-HCl, pH 7.4, 500 mM NaCl, 10% [v/v], 1% [v/v] Triton X-100, 1 mM EDTA) with no resuspension. Beads were then washed twice by resuspension in ice-cold 500 mM NaCl wash buffer followed by a final wash in ice-cold TBS (20 mM Tris-HCl, pH 7.4, 137 mM NaCl). Beads were resuspended in 15 μl TBS with 25 μl 5X sample loading buffer containing 2 mM biotin and incubated at 95°C for 10 mins. Beads and supernatants were stored at −20°C prior to SDS-PAGE and immunoblot analysis.

#### pTyr immunoprecipitation

400 μl dephosphorylation reactions (prepared as described above) were diluted to a total volume of 800 μl (0.2% [w/v] SDS final concentration) in 150 mM NaCl wash buffer (20 mM Tris-HCl, pH 7.4, 150 mM NaCl, 10% [v/v] glycerol, 1% [v/v] Triton X-100, 1 mM EDTA). 5 μl of rabbit anti-pTyr antibody (Cell Signalling Technology) was added to each sample and incubated for 3 h at 4 °C with rotation. Immunoprecipitation was carried out by addition of 40 μl of washed protein G agarose beads and samples were incubated overnight (16 h) at 4 °C with rotation. Beads were collected by centrifugation at 15000 xg for 30s at 4 °C and washed five times with 1 ml of ice-cold 150 mM wash buffer. After washing, beads were resuspended in 2.5X sample loading buffer and incubated at 95°C for 10 mins. Supernatants were collected and stored at −20 °C prior to SDS-PAGE and immunoblot analysis.

#### CRISPR/Cas9 genome editing

Oligonucleotides encoding single guide RNAs (sgRNAs) targeting human *PTPRU* exon 14 and exon 23 were cloned into pSpCas9.(BB).mCherry and pSpCas9.(BB).eGFP respectively as previously described (11). MCF10A cells were co-transfected with both plasmids by reverse transfection using Lipofectamine LTX/PLUS reagent as per manufacturer’s instructions. Clonal cell-lines were established by single-cell sorting of mCherry-eGFP double positive cells by flow cytometry, 48 h post-transfection. After expansion of clones, PTPRU-KO clones were identified using immunoblot for PTPRU. sgRNA target sites were amplified from genomic DNA to confirm editing. 3 independent confirmed PTPRU-KO clones were pooled to establish the final PTPRU-KO MCF10A population. PTPRK-KO MCF10A cells were generated in a previous study (1).

#### Protein sequence alignments

Protein multiple sequence alignments were generated using Clustal Omega (12) and edited using Jalview (13).

**Fig. S1.**
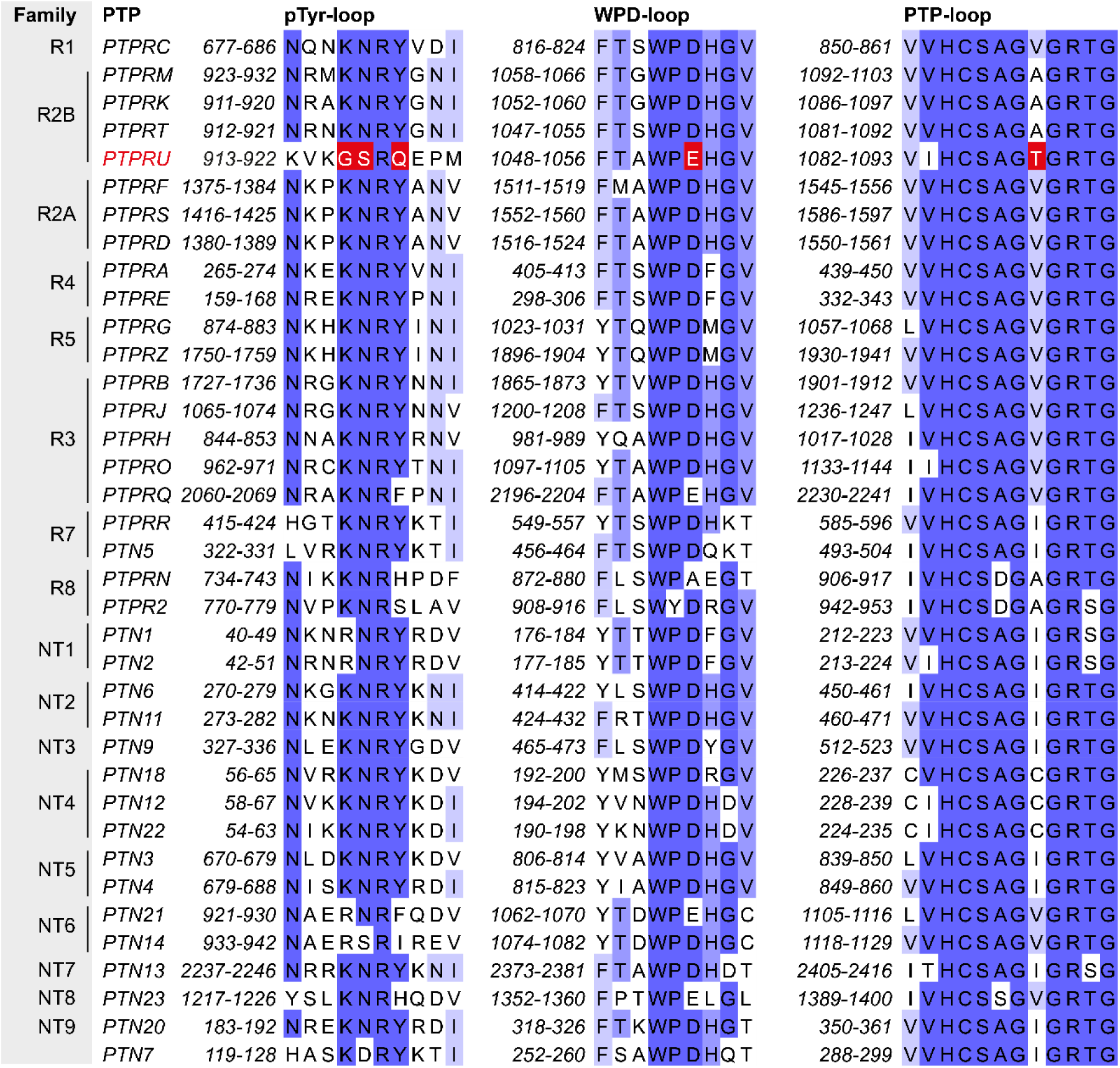
Multiple sequence alignment of the pTyr recognition loop, WPD loop and PTP loop of the 37 classical PTPs, colored by percentage identity (blue). Key variable residues in PTPRU are highlighted in red.

**Fig. S2.**
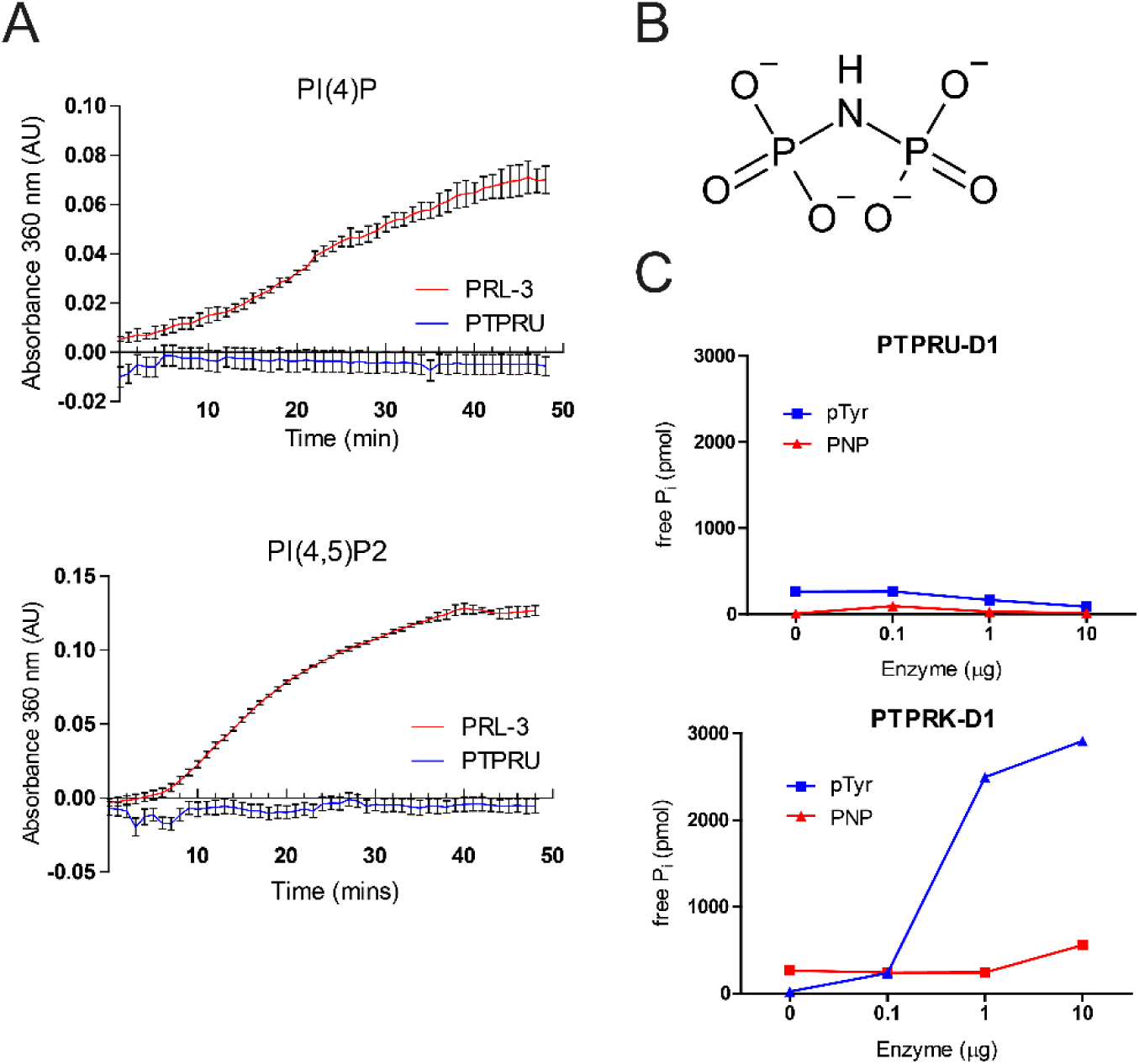
The PTPRU-D1 domain shows no activity against diverse substrates. *(A)* Phosphatidylinositol (PI) phosphatase activity assay. Phosphatidylinositol 4-phosphate [PI(4)P] and phosphatidyl 4,5-bisphosphate [PI(4,5)P2] substrates were incubated with either 3 μM PTPRU-D1 or 6 μM PRL-3 (positive control for PI phosphatase activity) (14) and product formation monitored by measurement of absorbance at 360 nm. *(B)* Chemical structure of the phosphoramidate-linked substrate imidodiphosphate (PNP). *(C)* Activity of PTPRU and PTPRK D1 domains incubated with 100 μM of either pTyr peptide or PNP, measured using BIOMOL Green reagent.

**Fig. S3.**
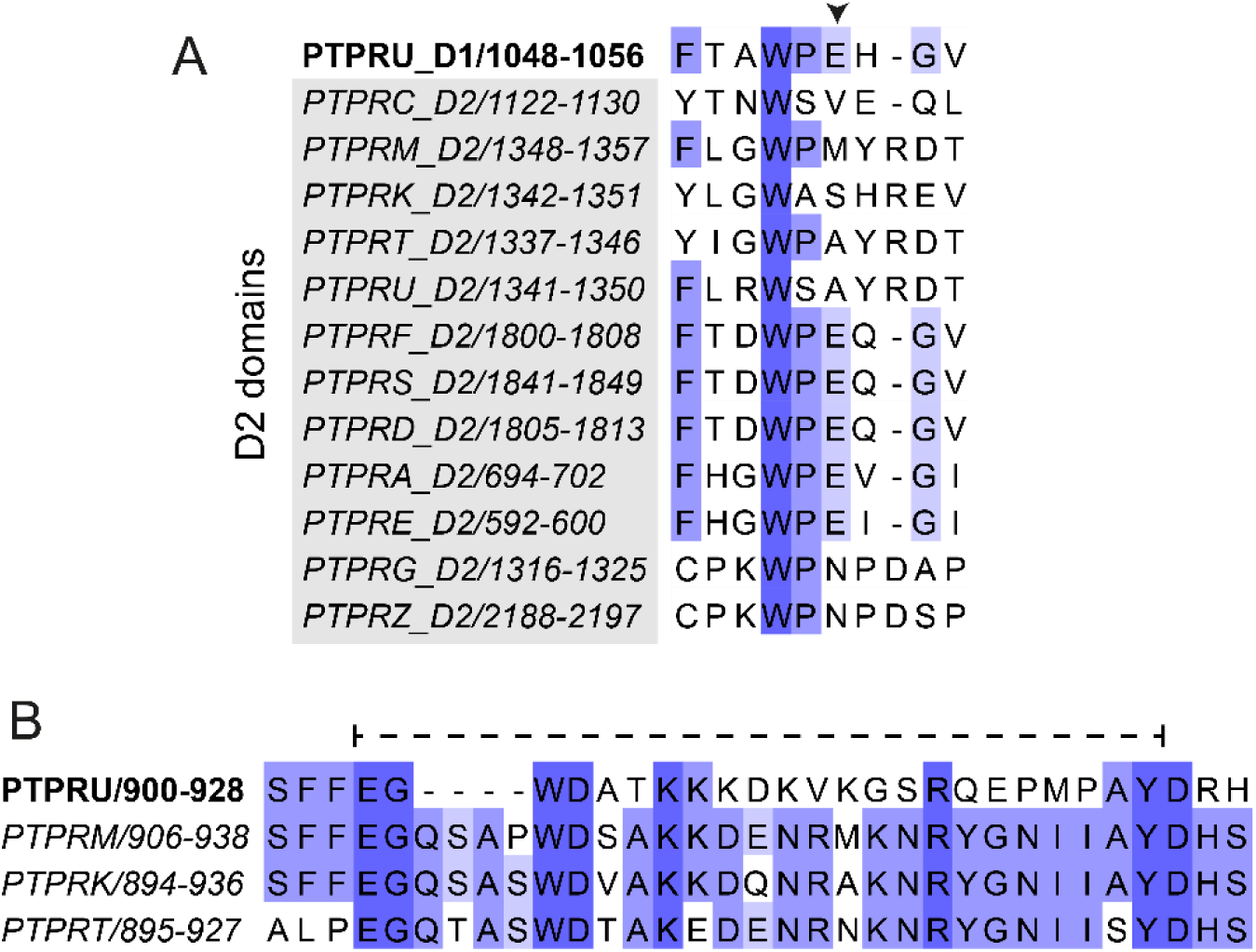
Sequence conservation of the PTPRU-D1 domain. *(A)* Multiple sequence alignment of the PTPRU-D1 WPD loop with those of the receptor PTP D2 domains, colored by percentage identity (blue). The non-canonical glutamate of the PTPRU-D1 WPD-loop is marked by an arrowhead. *(B)* Multiple sequence alignment of the R2B family pTyr recognition loops colored by percentage identity (blue). Residues 904-925 of PTPRU, which are disordered in the structure, are highlighted by a dashed line.

**Fig. S4.**
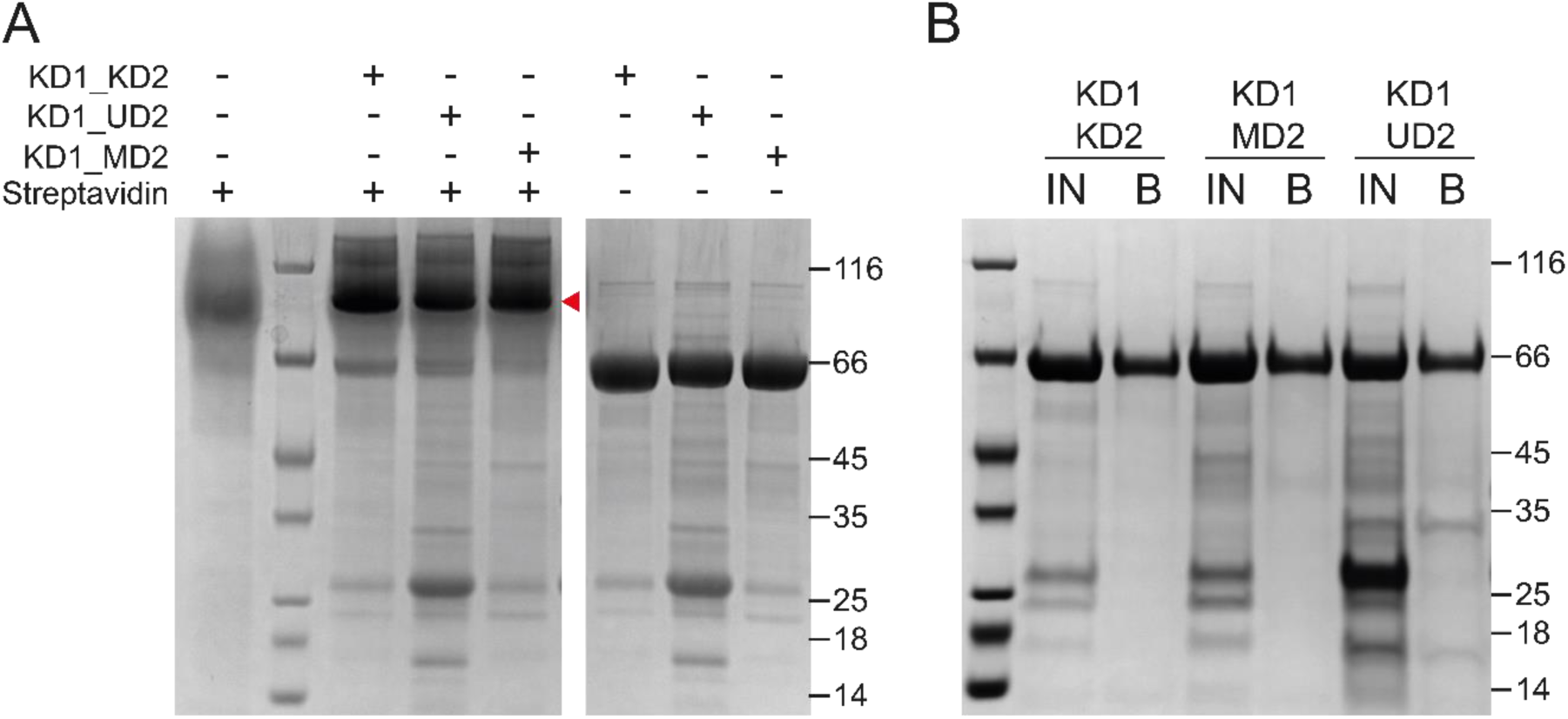
*In vivo* biotinylation of chimeric tandem PTP domains. *(A) In vivo* biotinylated chimeric tandem PTP domains incubated with or without streptavidin, resolved by SDS-PAGE and visualized by Coomasssie staining. Mobility shift upon streptavidin binding to biotinylated protein is indicated by arrowhead. *(B) In vivo* biotinylated chimeric tandem PTP domains bound to streptavidin magnetic beads. Input protein (IN) and protein eluted from washed beads (B) was resolved by SDS-PAGE and visualized by Coomassie staining.

**Table S1.** PTP domain structures used in structural alignments (Fig 3A)

| **PTP** | **PDB ID** | **RMSD (Å)\*** |
| --- | --- | --- |
| PTPN5 | 2BIJ | 1.020 over 1010 atoms |
| PTPN6 | 4HJP | 1.040 over 1082 atoms |
| PTPN11 | 3B7O | 1.015 over 1121 atoms |
| PTPN9 | 2PA5 | 0.946 over 1208 atoms |
| PTPN18 | 2OC3 | 1.248 over 1029 atoms |
| PTPN22 | 2P6X | 1.172 over 1085 atoms |
| PTPN3 | 2B49 | 1.010 over 1139 atoms |
| PTPN4 | 2I75 | 1.063 over 1121 atoms |
| PTPN14 | 2BZL | 1.104 over 1102 atoms |
| PTPN13 | 1WCH | 1.042 over 1038 atoms |
| PTPRC | 1YGU | 1.082 over 1255 atoms |
| PTPRF | 1LAR | 0.811 over 1233 atoms |
| PTPRS | 2FH7 | 0.864 over 1446 atoms |
| PTPRM | 1RPM | 0.788 over 1505 atoms |
| PTPRK | 2C7S | 0.804 over 1582 atoms |
| PTPRT | 2OOQ | 0.874 over 1500 atoms |
| PTPRB | 2AHS | 0.989 over 1337 atoms |
| PTPRJ | 2NZ6 | 0.954 over 1216 atoms |
| PTPRO | 2GJT | 1.081 over 1332 atoms |
| PTPRA | 1YFO | 0.881 over 1334 atoms |
| PTPRE | 2JJD | 0.865 over 1204 atoms |
| PTPRG | 2H4V | 0.989 over 1416 atoms |
| PTPRR | 2A8B | 1.162 over 1170 atoms |
| PTPN7 | 2A3K | 0.962 over 1095 atoms |
| PTPRN | 2I1Y | 1.073 over 1142 atoms |
| PTPRN2 | 2QEP | 0.895 over 1062 atoms |
| PTPN1 | 2NT7 | 1.223 over 1116 atoms |

\*Calculated using extra\_fit function within Pymol using default settings (5 cycles, cut-off = 2.0 Å) with PTPRU-D1 (reduced) as the target molecule.

**Table S2.** Percentage sequence identity matrix of R2B family D1 domains vs R2B family D1 and D2 domains. Generated by multiple sequence alignment using Clustal Omega.

|  | PTPRU<br>D1 | PTPRK<br>D1 | PTPRM<br>D1 | PTPRT<br>D1 | PTPRU<br>D2 | PTPRK<br>D2 | PTPRM<br>D2 | PTPRT<br>D2 |
| --- | --- | --- | --- | --- | --- | --- | --- | --- |
| PTPRU<br>D1 | 100 | 72.44 | 69.61 | 64.66 | 27.34 | 29.2 | 28.15 | 26.64 |
| PTPRK<br>D1 | 72.44 | 100 | 79.44 | 76.31 | 28.37 | 31.29 | 31.02 | 28.78 |
| PTPRM<br>D1 | 69.61 | 79.44 | 100 | 80.14 | 27.66 | 30.58 | 30.29 | 27.34 |
| PTPRTD<br>1 | 64.66 | 76.31 | 80.14 | 100 | 29.79 | 30.58 | 30.29 | 29.14 |

## REFERENCES

1. Elchebly M, et al. (1999) Increased insulin sensitivity and obesity resistance in mice lacking the protein tyrosine phosphatase-1B gene. Science 283(5407):1544–1548.

2. Byth KF, et al. (1996) CD45-null transgenic mice reveal a positive regulatory role for CD45 in early thymocyte development, in the selection of CD4+CD8+ thymocytes, and B cell maturation. J Exp Med 183(4):1707–1718.

3. Fearnley GW, et al. (2019) The homophilic receptor PTPRK selectively dephosphorylates multiple junctional regulators to promote cell–cell adhesion. eLife 8:e44597.

4. Chen Y-NP, et al. (2016) Allosteric inhibition of SHP2 phosphatase inhibits cancers driven by receptor tyrosine kinases. Nature 535:148.

5. Krishnan N, et al. (2014) Targeting the disordered C terminus of PTP1B with an allosteric inhibitor. Nat Chem Biol 10(7):558–566.

6. Guan KL & Dixon JE (1991) Evidence for protein-tyrosine-phosphatase catalysis proceeding via a cysteine-phosphate intermediate. J Biol Chem 266(26):17026–17030.

7. Andersen JN, et al. (2001) Structural and Evolutionary Relationships among Protein Tyrosine Phosphatase Domains. Mol Cell Biol 21(21):7117–7136.

8. Juettner VV, et al. (2019) VE-PTP stabilizes VE-cadherin junctions and the endothelial barrier via a phosphatase-independent mechanism. J Cell Biol 218(5):1725–1742.

9. Stewart RA, et al. (2010) Phosphatase-dependent and -independent functions of Shp2 in neural crest cells underlie LEOPARD syndrome pathogenesis. Dev Cell 18(5):750–762.

10. Coughlin S, et al. (2015) An extracatalytic function of CD45 in B cells is mediated by CD22. Proc Natl Acad Sci U S A 112(47):E6515–6524.

11. Barr AJ, et al. (2009) Large-Scale Structural Analysis of the Classical Human Protein Tyrosine Phosphatome. Cell 136(2):352–363.

12. Gingras MC, et al. (2009) HD-PTP is a catalytically inactive tyrosine phosphatase due to a conserved divergence in its phosphatase domain. PLoS One 4(4):e5105.

13. Chen K-E, et al. (2015) Substrate Specificity and Plasticity of FERM-Containing Protein Tyrosine Phosphatases. Structure 23(4):653–664.

14. Magistrelli G, Toma S, & Isacchi A (1996) Substitution of two variant residues in the protein tyrosine phosphatase-like PTP35/IA-2 sequence reconstitutes catalytic activity. Biochem Biophys Res Commun 227(2):581–588.

15. Gingras MC, et al. (2009) Expression analysis and essential role of the putative tyrosine phosphatase His-domain-containing protein tyrosine phosphatase (HD-PTP). Int J Dev Biol 53(7):1069–1074.

16. Bend R, et al. (2019) Phenotype and mutation expansion of the PTPN23 associated disorder characterized by neurodevelopmental delay and structural brain abnormalities. Eur J Hum Genet.

17. van der Wijk T, Overvoorde J, & den Hertog J (2004) H2O2-induced Intermolecular Disulfide Bond Formation between Receptor Protein-tyrosine Phosphatases. J Biol Chem 279(43):44355–44361.

18. Toledano-Katchalski H, et al. (2003) Dimerization In Vivo and Inhibition of the Nonreceptor Form of Protein Tyrosine Phosphatase Epsilon. Mol Cell Biol 23(15):5460.

19. Oganesian A, et al. (2003) Protein tyrosine phosphatase RQ is a phosphatidylinositol phosphatase that can regulate cell survival and proliferation. Proc Natl Acad Sci U S A 100(13):7563–7568.

20. Sommer L, Rao M, & Anderson DJ (1997) RPTPδ and the novel protein tyrosine phosphatase RPTPψ are expressed in restricted regions of the developing central nervous system. Developmental Dyn 208(1):48–61.

21. Aerne B, Stoker A, & Ish-Horowicz D (2003) Chick receptor tyrosine phosphatase Ψ is dynamically expressed during somitogenesis. Gene Expr Patterns 3(3):325–329.

22. Aerne B & Ish-Horowicz D (2004) Receptor tyrosine phosphatase psi is required for Delta/Notch signalling and cyclic gene expression in the presomitic mesoderm. Development 131(14):3391–3399.

23. Badde A & Schulte D (2008) A Role for Receptor Protein Tyrosine Phosphatase λ in Midbrain Development. J Neurosci. 28(24):6152.

24. Liu Y, et al. (2014) Knockdown of protein tyrosine phosphatase receptor U inhibits growth and motility of gastric cancer cells. Int J Clin Exp Pathol. 7(9):5750–5761.

25. Wang H, et al. (2014) Protein tyrosine phosphatase receptor U (PTPRU) is required for glioma growth and motility. Carcinogenesis 35(8):1901–1910.

26. Becka S, et al. (2010) Characterization of the adhesive properties of the type IIb subfamily receptor protein tyrosine phosphatases. Cell Commun Adhes 17(2):34–47.

27. Yan H-X, et al. (2002) Physical and Functional Interaction between Receptor-like Protein Tyrosine Phosphatase PCP-2 and β-Catenin. Biochemistry 41(52):15854–15860.

28. Yan H-X, et al. (2006) Protein-tyrosine Phosphatase PCP-2 Inhibits β-Catenin Signaling and Increases E-cadherin-dependent Cell Adhesion. J Biol Chem 281(22):15423–15433.

29. Kumar P, et al. (2017) A Human Tyrosine Phosphatase Interactome Mapped by Proteomic Profiling. J Proteome Res 16(8):2789–2801.

30. Sarmiento M, Zhao Y, Gordon SJ, & Zhang ZY (1998) Molecular basis for substrate specificity of protein-tyrosine phosphatase 1B. J Biol Chem 273(41):26368–26374.

31. Hardman G, et al. (2019) Strong anion exchange-mediated phosphoproteomics reveals extensive human non-canonical phosphorylation. EMBO J:e100847.

32. Yokoi F, Hiraishi H, & Izuhara K (2003) Molecular cloning of a cDNA for the human phospholysine phosphohistidine inorganic pyrophosphate phosphatase. J Biochem 133(5):607–614.

33. Srivastava S, et al. (2008) Protein histidine phosphatase 1 negatively regulates CD4 T cells by inhibiting the K+ channel KCa3.1. Proc Natl Acad Sci U S A 105(38):14442–14446.

34. Panda S, et al. (2016) Identification of PGAM5 as a Mammalian Protein Histidine Phosphatase that Plays a Central Role to Negatively Regulate CD4(+) T Cells. Mol Cell 63(3):457–469.

35. Hiraishi H, Yokoi F, & Kumon A (1998) 3-phosphohistidine and 6-phospholysine are substrates of a 56-kDa inorganic pyrophosphatase from bovine liver. Arch Biochem Biophys 349(2):381–387.

36. Flint AJ, Tiganis T, Barford D, & Tonks NK (1997) Development of “substrate-trapping” mutants to identify physiological substrates of protein tyrosine phosphatases. Proc Natl Acad Sci U S A 94(5):1680–1685.

37. Eswaran J, Debreczeni JE, Longman E, Barr AJ, & Knapp S (2006) The crystal structure of human receptor protein tyrosine phosphatase kappa phosphatase domain 1. Protein Sci 15(6):1500–1505.

38. Salmeen A, Andersen JN, Myers MP, Tonks NK, & Barford D (2000) Molecular basis for the dephosphorylation of the activation segment of the insulin receptor by protein tyrosine phosphatase 1B. Mol Cell 6(6):1401–1412.

39. Machado L, Critton DA, Page R, & Peti W (2017) Redox Regulation of a Gain-of-Function Mutation (N308D) in SHP2 Noonan Syndrome. ACS Omega 2(11):8313–8318.

40. Tsai SJ, et al. (2009) Crystal structure of the human lymphoid tyrosine phosphatase catalytic domain: insights into redox regulation. Biochemistry 48(22):4838–4845.

41. Lee CU, et al. (2015) Redox Modulation of PTEN Phosphatase Activity by Hydrogen Peroxide and Bisperoxidovanadium Complexes. Angew Chem Int Ed Engl 54(46):13796–13800.

42. Machado LESF, Shen T-L, Page R, & Peti W (2017) The KIM-family protein-tyrosine phosphatases use distinct reversible oxidation intermediates: Intramolecular or intermolecular disulfide bond formation. J Biol Chem 292(21):8786–8796.

43. van der Wijk T, Blanchetot C, Overvoorde J, & den Hertog J (2003) Redox-regulated Rotational Coupling of Receptor Protein-tyrosine Phosphatase α Dimers. J Biol Chem 278(16):13968–13974.

44. van Montfort RL, Congreve M, Tisi D, Carr R, & Jhoti H (2003) Oxidation state of the active-site cysteine in protein tyrosine phosphatase 1B. Nature 423(6941):773–777.

45. Salmeen A, et al. (2003) Redox regulation of protein tyrosine phosphatase 1B involves a sulphenyl-amide intermediate. Nature 423:769.

46. Evans PR & Murshudov GN (2013) How good are my data and what is the resolution? Acta Crystallogr D Biol Crystallogr 69(Pt 7):1204–1214.

47. Emsley P, Lohkamp B, Scott WG, & Cowtan K (2010) Features and development of Coot. Acta Crystallogr D Biol Crystallogr 66(Pt 4):486–501.

48. Afonine PV, et al. (2012) Towards automated crystallographic structure refinement with phenix.refine. Acta Crystallogr D Biol Crystallogr 68(Pt 4):352–367.

## SI References

1. Fearnley GW, et al. (2019) The homophilic receptor PTPRK selectively dephosphorylates multiple junctional regulators to promote cell–cell adhesion. eLife 8:e44597.

2. Becka S, et al. (2010) Characterization of the adhesive properties of the type IIb subfamily receptor protein tyrosine phosphatases. Cell Commun Adhes 17(2):34–47.

3. Winter G, et al. (2018) DIALS: implementation and evaluation of a new integration package. Acta Crystallogr D Struct Biol 74(Pt 2):85–97.

4. Evans PR & Murshudov GN (2013) How good are my data and what is the resolution? Acta Crystallogr D Biol Crystallogr 69(Pt 7):1204–1214.

5. McCoy AJ, et al. (2007) Phaser crystallographic software. J Appl Crystallogr 40(Pt 4):658–674.

6. Eswaran J, Debreczeni JE, Longman E, Barr AJ, & Knapp S (2006) The crystal structure of human receptor protein tyrosine phosphatase kappa phosphatase domain 1. Protein Sci 15(6):1500–1505.

7. Emsley P, Lohkamp B, Scott WG, & Cowtan K (2010) Features and development of Coot. Acta Crystallogr D Biol Crystallogr 66(Pt 4):486–501.

8. Afonine PV, et al. (2012) Towards automated crystallographic structure refinement with phenix.refine. Acta Crystallogr D Biol Crystallogr 68(Pt 4):352–367.

9. Soule HD, et al. (1990) Isolation and characterization of a spontaneously immortalized human breast epithelial cell line, MCF-10. Cancer Res 50(18):6075–6086.

10. Schindelin J, et al. (2012) Fiji - an Open Source platform for biological image analysis. Nat Methods 9(7):10.1038/nmeth.2019.

11. Ran FA, et al. (2013) Genome engineering using the CRISPR-Cas9 system. Nat Protoc 8:2281.

12. Madeira F, et al. (2019) The EMBL-EBI search and sequence analysis tools APIs in 2019. Nucleic Acids Res 47(W1):W636–W641.

13. Waterhouse AM, Procter JB, Martin DM, Clamp M, & Barton GJ (2009) Jalview Version 2--a multiple sequence alignment editor and analysis workbench. Bioinformatics 25(9):1189–1191.

14. McParland V, et al. (2011) The metastasis-promoting phosphatase PRL-3 shows activity toward phosphoinositides. Biochemistry 50(35):7579–7590.

